# Investigating the necessity of neuronal and oligodendroglial mutant ATXN3 expression for Spinocerebellar ataxia type 3 phenotypes

**DOI:** 10.64898/2026.11.23.768600

**Authors:** Alexandra F. Putka, Jacen Emerson, Brianna S. Nelthrope, Vivian I. Schmidt, Benjamin D. Carr, Toluwani T. Adelaja, Grace Mao, Juan P. Mato, Addison R. Kim, Hannah K. Shorrock, Hayley S. McLoughlin

**Author notes:** These authors contributed equally to this work (Alexandra F. Putka, Jacen Emerson).

## Abstract

There is growing evidence for cell type-specific vulnerability across the neurodegenerative disease field, and Spinocerebellar ataxia type 3 (SCA3) is no exception. SCA3 is caused by a CAG repeat expansion in the *ATXN3* gene, leading to progressive motor incoordination and premature mortality. Unfortunately, this debilitating disease has no FDA-approved treatments, necessitating ongoing work to identify underlying drivers of pathogenesis. While neurons and oligodendrocytes are well-established as vulnerable cell types in SCA3, the causal contribution of these cells to disease phenotypes remained unknown. In this study, we harnessed a novel conditional Knock-in mouse model of SCA3 wherein the mutant *Atxn3* gene can be selectively silenced in any cell expressing Cre. With this mouse we interrogated the necessity of mutant *Atxn3* expression in neurons constitutively targeted by *Baf53b-Cre*, oligodendrocytes and Schwann cells inducibly targeted by *Sox10-iCre/ERT2*, and dual targeted neural crest-derived cells and oligodendrocytes by *Sox10-Cre*. We found limited attenuation of SCA3 motor deficits when mutant ATXN3 was silenced in neurons or oligodendrocytes/Schwann cells alone. Rather, targeting oligodendrocytes and the neural crest-derived lineage produced partial rescue of total locomotion and full rescue of balance beam traversal latency. Bulk RNA sequencing and proteomics on the disease-vulnerable cerebellum of conditional SCA3 mice revealed that oligodendrocyte maturation did not correspond with behavioral rescue, but inflammatory processes and alternative splicing did. This work suggests that synergistic interactions between neurons and oligodendrocytes contribute to SCA3 behavioral and neuropathological phenotypes and highlights the requirement of targeting multiple cell types for therapeutic benefit.

## Introduction

Spinocerebellar ataxia type 3 (SCA3) is a debilitating and ultimately fatal neurodegenerative disease caused by a CAG repeat expansion in the *ATXN3* gene. Patients with SCA3 experience progressive impairments in balance, coordination, swallowing, and speaking, uniformly leading to death 10 to 20 years after disease onset [1–4]. Although SCA3 is the most common dominantly inherited ataxia, no FDA-approved treatments exist. In part, this is because of a gap in knowledge of the cellular and molecular mechanisms linking the CAG repeat expansion in *ATXN3* with disease symptoms. Interestingly, despite *ATXN3* expression across all major organ systems of the body, SCA3 pathology is restricted to select vulnerable regions in the nervous system. Vulnerable regions include the deep cerebellar nuclei, pontine nuclei, and spinal cord [2,3], along with more recently described regions such as the dorsal root ganglia (DRG) and peripheral nerves [5–7]. Beyond regional vulnerability, there is growing evidence of cell type-specific vulnerabilities in SCA3, though the causal relationship of specific cell types to SCA3 pathogenesis remains understudied.

Similar to many other neurodegenerative diseases, neurons are the most well-described of the affected cell types in SCA3. Indeed, hallmark nuclear ATXN3 accumulation was first discovered in this cell type [8,9], with additional pathological processes including perturbed protein quality control, mitochondrial function, synaptic function, and electrophysiology [2,10–12]. Beyond neurons, recent work shows dysfunction in oligodendrocytes, the myelinating cells of the central nervous system. Our lab demonstrated an early reduction in oligodendrocyte maturation in SCA3-vulnerable brain regions of mouse models, corresponding with myelin abnormalities in SCA3 patients and mice [13–16]. As for shared mechanisms of dysfunction in neurons and oligodendrocytes, recent work from our lab uncovered broad RNA splicing dysregulation in SCA3 mice, particularly affecting genes related to synaptic function and oligodendrocyte maturation [17,18]. While there is strong evidence that neuron and oligodendrocyte dysfunction is correlated with SCA3 disease phenotypes, there is a notable dearth of research on the causal role of these cells.

Prior studies addressing cell type-specific contributions to SCA3 provide insight into pathogenic mechanisms, but methodological limitations leave important questions unresolved. For example, SCA3 cortical neurons derived from human embryonic stem cells demonstrate cell-autonomous protein dyshomeostasis [19], while primary oligodendrocyte precursor cells from SCA3 mice recapitulate disease-associated maturation impairments [13,14]. However, *in vitro* cell models cannot clarify how mutant ATXN3 expression in specific cell types contributes to SCA3-associated behavioral phenotypes. One study sought to answer this question *in vivo* using a transgenic mouse model expressing truncated human mutant *ATXN3* cDNA specifically in cerebellar Purkinje cells [20]. These mice developed an early and severe ataxic phenotype, suggesting that overexpression of mutant ATXN3 in Purkinje cells is sufficient to drive SCA3-associated motor impairments. However, this model restricts mutant ATXN3 overexpression to a single neuronal population and therefore does not recapitulate the broad neuronal dysfunction induced by endogenous levels of mutant ATXN3 across SCA3-vulnerable brain regions [3]. Together, these studies emphasize the need for an *in vivo* tool enabling cell type-specific studies of endogenous mutant ATXN3.

Our lab recently developed a conditional Knock-in (cKI) mouse model of SCA3, in which mutant ATXN3 can be selectively silenced in any cell type expressing Cre recombinase [7]. This model avoids the confounding effects of transgene overexpression. Consistent with key features of human disease, these mice develop motor deficits as early as 12 weeks of age, with progressive impairments documented through 36 weeks [7]. Thus, this cKI model provides a novel tool for defining the causal contributions of specific cell types to SCA3-associated phenotypes.

In the present study, we evaluated whether mutant ATXN3 expression in specific cell types is necessary for SCA3 behavioral, transcriptomic, and proteomic phenotypes. Mutant ATXN3 was silenced either constitutively in neurons using *Baf53b-Cre*, inducibly in oligodendrocytes and Schwann cells using *Sox10-iCre/ERT2*, or constitutively in both populations using the neural crest-lineage driver *Sox10-Cre*. Silencing mutant ATXN3 in either neurons or oligodendrocytes/Schwann cells alone failed to produce sustained rescue of SCA3 motor phenotypes, whereas targeting both populations led to synergistic and sustained rescue. Bulk RNA sequencing and proteomic analysis of the cerebellum in late-stage disease revealed partial rescue of neuroinflammatory signatures but not oligodendrocyte maturation deficits. Consistent with previous studies from our lab, RNA splicing abnormalities were prominent in the cerebellum and were substantially rescued by cell type-specific mutant ATXN3 silencing. Overall, these findings define the contributions of specific cell populations to SCA3-associated disease signatures and emphasize the importance of targeting multiple cell types for therapeutic benefit.

## Materials and methods

### Mouse models

All animal procedures were approved by the University of Michigan Institutional Animal Care and Use Committee and conducted in accordance with the United States Public Health Service’s policy on Humane Care and Use of Laboratory Animals. Mice were housed in a room with standard 12-hour light/dark cycles and food and water provided *ad libitum*. The conditional Knock-In *Atxn3^cQ300/Q6^* (cKI) mice utilized in this study harbor one *Atxn3* allele with a hyperexpanded CAG_300_ tract and loxP sites flanking exon 4 [7]. The remaining allele of *Atxn3* retains the endogenous CAG_6_ tract and has no loxP sites. As previously described, Cre-mediated recombination of the loxP sites around exon 4 causes a change in amino acid sequence at residue 78, leading to early truncation of ATXN3 10 amino acids later [7]. cKI mice were genotyped using tail biopsy DNA collected before weaning and confirmed postmortem, as previously described [7]. For selective silencing of the mutant *Atxn3^cQ300^* allele in neurons, cKI mice were crossed with the constitutive driver *Baf53b-Cre* (JAX #027826) [21]. To target myelinating glia (oligodendrocytes and Schwann cells), cKI mice were crossed with inducible *Sox10-iCreERT2* (JAX #027651) mice [22]. To target neural crest-derived cells and the oligodendrocyte lineage, cKI mice were crossed with the constitutive driver *Sox10-Cre* (JAX #025807) [23]. For reporter studies, cKI; Cre strains were crossed to Ai14 tdTomato reporter mice (JAX #007914) [24]. Cre and tdTomato mice were genotyped according to Jackson Laboratory protocols. All strains were maintained on a C57BL/6J background. MiniMUGA Single Nucleotide Polymorphism (SNP) panel analysis (Transnetyx) demonstrated the percent of diagnostic SNPs explained by the C57BL/6J background was: 90.0% for cKI, 88.5% for *Baf53b-Cre*, 97.6% in *Sox10-iCre/ERT2*, and 83.6% for *Sox10-Cre* mice. At collection timepoints, ketamine/xylazine-anesthetized mice were perfused with 1X Phosphate-buffered saline (PBS). The left hemisphere cerebellum and frontal cortex were flash frozen for biochemical experiments. The right hemisphere of the brain, along with the spinal cord and dorsal root ganglia (DRG), were post-fixed in 4% paraformaldehyde (PFA) overnight for immunofluorescence experiments.

#### Tamoxifen treatment

For Cre induction in *Sox10-iCreERT2* mice only, tamoxifen (Sigma-Aldrich, T5648) was dissolved in corn oil (Sigma-Aldrich, C8267) at 20 mg/mL by overnight shaking incubation at 37°C [25]. At 12 weeks of age (mean: 12.11 weeks, SEM: 0.027), *Sox10-iCreERT2*+ cKI and WT mice were injected intraperitoneally with tamoxifen (100 mg/kg) or vehicle once per day for five consecutive days [25–28]. Each cohort of mice received a fresh preparation of tamoxifen in corn oil, which was stored at 4°C for the duration of injections.

#### Mouse behavior

Mouse motor behavior was assessed longitudinally at 12, 24, and 36 weeks of age in *Baf53b-Cre* and *Sox10-Cre* mice. Data from cKI; Cre+ mice of each strain were compared to sex- and age-matched WT; Cre- and cKI; Cre- controls. In *Sox10-iCreERT2* mice, motor behavior was assessed longitudinally at 11, 24, and 36 weeks, with 11 weeks being one week prior to tamoxifen or vehicle treatment. Data from cKI; *iSox10*+; tamoxifen mice were compared to sex- and age-matched cKI; *iSox10*+; Vehicle and WT; *iSox10*+; Vehicle controls. At each timepoint, mouse weight was recorded prior to balance beam and locomotor activity assessments, as previously described [7,12,16]. Balance and coordination were assessed by measuring the latency to traverse a 5-mm round beam toward a dark box filled with mouse bedding. Mice were exposed to two consecutive days of training before recording data on the third day, with the traversal latency averaged across two trials per mouse. Falls from the beam were recorded as 30 seconds. Total locomotor activity (x/y-axis beam breaks) was recorded over a 30-minute trial by placing mice in a photobeam activity system open field apparatus (San Diego Instruments).

### RNA extraction and expression analysis

Flash frozen cerebella collected from 36-week-old WT; Cre-, cKI; Cre-, and cKI; Cre+ mice of both the *Baf53b-Cre* and *Sox10-Cre* strains (n =3–4 mice/genotype/strain) were homogenized in RIPA buffer (Sigma-Aldrich, R0278) with protease inhibitors (Sigma-Aldrich, 11836170001) and the lysate was aliquoted for either RNA or protein extraction. Lysate for RNA extraction was combined with TRIzol reagent (Invitrogen, 15596018) and processed, as previously described [12,16]. RNA quality was assessed using the DNF-471 RNA kit (Agilent) on a 5300 fragment analyzer (Agilent Bioanalyzer) and concentration quantified using the Qubit RNA high sensitivity assay (Thermo Fisher Scientific). Libraries were prepared from 200ng RNA using rRNA Depletion (NEB, E7405) and the NEBNext Ultra II Directional RNA Library Preparation Kit for Illumina (NEB, E7765). Library quality was assessed via the DNF-474 HS NGS kit (Agilent) and quantification performed using NEBNext Library Quantification Kit for Illumina (NEB, E7630). Equivalent library quantities were loaded on a P4 Flowcell (Illumina, 20100992) and run on an Illumina NextSeq 2000. The resulting FASTQ files were quality checked using FastQC, and all samples with at least 30 million paired end reads were included in the study. All Fastq files were trimmed using fastp (version 1.3.6) and some were downsampled using seqtk (version 1.5-r133) to ensure comparable library sizes. Per sample processing details can be found in **Supplementary Table 1**. Processed fastq files were then aligned to the GRCm39 mouse reference genome using STAR (version 2.7.10b) [29]. Differential gene expression analysis was performed using DESeq2 (version 1.46.0) [30] using thresholds of p-adjusted < 0.05 and |log2FC| > 0.5849 (a fold-change of 1.5). Phenotypic gene changes were significant by p-adjusted < 0.05 and |log2FC| > 0.5849 in the cKI; Cre- vs. WT; Cre- comparison. Cre-driven gene changes were defined from a list of genes not significantly altered between cKI; Cre- and WT; Cre- mice by either the p-adjusted or log2FC cutoff that were significantly altered in both cKI; Cre+ vs. cKI; Cre- mice and cKI; Cre+ vs. WT; Cre- mice. Rescue was defined as a gene with a log2FC < -0.152 or log2FC > 0.138 (a 10% change) in cKI; Cre+ mice relative to cKI; Cre- mice. Functional enrichment analysis was conducted with g:Profiler, with results shown for Gene Ontology (GO) terms [31,32].

Alternative splicing analysis was carried out in rMATS (version 4.1.2) [33] with significance thresholds of FDR < 0.05 and ΔPSI > 10%. Phenotypic splicing events were significant by FDR < 0.05 and ΔPSI > 10% in the cKI; Cre- vs. WT; Cre- comparison. Cre-driven splicing events were defined from a list of events not significantly altered between cKI; Cre- and WT; Cre- mice by either the FDR or the ΔPSI cutoff that were significantly altered in both cKI; Cre+ vs. cKI; Cre- mice and cKI; Cre+ vs. WT; Cre- mice. Rescue was defined as an average PSI in cKI; Cre+ mice at least 10% different from cKI; Cre- mice, in the direction of WT; Cre- controls.

### Quantitative PCR (qPCR)

Flash-frozen cerebella from WT; *Sox10-Cre-*, cKI; *Sox10-Cre-*, and cKI; *Sox10-Cre+* mice at 12 and 24 weeks of age underwent RNA extraction, as described above. Following Nanodrop to measure RNA concentration and purity, RNA was converted into cDNA using the iScript cDNA Synthesis Kit (Bio-Rad, 1708890) according to manufacturer recommendations. The resultant cDNA was diluted 1:10 in nuclease-free water. Finally, qPCR was conducted using mouse TaqMan primers, multiplexed for the transcript of interest (FAM) and *Beta-actin* (VIC), and run on a QuantStudio 3 qPCR machine (Thermo Fisher Scientific) under the TaqMan Fast Protocol [15]. TaqMan primers from Thermo Fisher Scientific include *Beta-actin* (Mm02619580_g1) and mature oligodendrocyte markers *Plp1* (Mm01297210_m1), *Mobp* (Mm02745649_m1), *Mal* (Mm01339780_m1), and *Ugt8a* (Mm00495930_m1). Relative quantification (RQ) of gene expression for each mature oligodendrocyte marker was calculated relative to WT; *Sox10-Cre-* mice at each timepoint and normalized to *Beta-actin*. Graphs represent RQ = 2^-(ddCT)^.

### Proteomics

#### Protein Digestion and TMT labeling

Cerebellar lysate from 36-week-old *Baf53b-Cre* and *Sox10-Cre* mice was processed for protein extraction, as previously described, with n =4–6 mice/genotype/strain [15]. Protein concentration was measured via a BCA assay (Thermo Fisher Scientific, 23227), and 75µg of protein (2µg/µL concentration) was submitted to the University of Michigan Proteomics Resource Facility. *Baf53b-Cre* and *Sox10-Cre* samples were run and analyzed separately. Samples were proteolyzed and isotopically labeled using the Tandem Mass Tag (TMT) 16-plex isobaric labeling reagent system (Thermo Fisher Scientific, A44521), following the manufacturer’s protocol with minor modifications. Briefly, cysteine residues were reduced and alkylated prior to protein precipitation by the addition of six volumes of ice-cold acetone and overnight incubation at −20°C. Precipitated protein was collected by centrifugation, and the resulting pellet was air-dried and resuspended in 100 µL of 0.1 M triethylammonium bicarbonate (TEAB). Proteins were digested overnight at 37°C with trypsin at a 1:40 enzyme-to-protein ratio, with continuous agitation on a thermomixer. TMT 16-plex reagents were reconstituted in 20 µL anhydrous acetonitrile, and labeling was carried out by transferring the complete digest into the TMT reagent vial and incubating at room temperature for 1 hour. The labeling reaction was quenched by the addition of 8 µL of 5% hydroxylamine followed by a 15-minute incubation. All labeled samples were then pooled and dried by vacuum centrifugation (vacufuge). The combined sample (∼200 μg) was fractionated offline into 8 fractions using a high-pH reversed-phase peptide fractionation kit (Thermo Fisher Scientific, 84868) per manufacturer instructions. Fractions were dried and reconstituted in 9 µL of 0.1% formic acid / 2% acetonitrile prior to LC-MS/MS analysis.

#### Mass Spectrometry Data Acquisition

To achieve accurate quantification, data were acquired using the multinotch-MS3 approach, which reduces reporter ion ratio distortion caused by co-isolated peptide interference during MS analysis [34]. All experiments were performed on an Orbitrap Ascend Tribrid mass spectrometer equipped with a FAIMS Pro source (Thermo Fisher Scientific) coupled to a Vanquish Neo UHPLC system. For each injection, 2 µL of sample was loaded onto an Easy-Spray PepMap Neo column (Thermo Fisher Scientific, 75 µm i.d. × 50 cm) and separated at a flow rate of 300 nL/minute using a binary gradient of 0.1% formic acid in water (solvent A) and 0.1% formic acid in acetonitrile (solvent B) as follows: 3–19% B over 72 minutes, 19–29% B over 28 minutes, 29–41% B over 20 minutes, followed by a 10-minute column wash at 95% B and re-equilibration to initial conditions. Eluting peptides were introduced into the mass spectrometer via an EasySpray ion source (Thermo Fisher Scientific). The FAIMS source was operated in standard resolution mode with a nitrogen gas flow of 4.2 L/minute, inner and outer electrode temperatures of 100°C, and a dispersion voltage of −5000 V. Ion selection was performed using two alternating compensation voltages (CVs) of −45 V and −65 V, with 1.5 seconds allocated per CV. The mass spectrometer was operated in a data-dependent acquisition mode. Full MS1 scans were acquired in the Orbitrap over a range of 400– 1600 m/z at a resolution of 120,000 (AGC target: 100%; maximum injection time: Auto). Precursor ions with charge states of +2 to +6 were isolated using the quadrupole mass filter at a 0.7 m/z isolation window and fragmented by collision-induced dissociation (CID) in the ion trap (NCE: 30%; normalized AGC target: 100%; maximum injection time: 35 ms). Real-Time Search (RTS) was performed against a mouse SwissProt protein database to identify and select TMT-labeled peptides for MS3 analysis in real time. For multinotch-MS3, the top 10 fragment ions from each RTS-qualified MS2 spectrum were selected and further fragmented by higher-energy collisional dissociation (HCD), with fragment ion detection in the Orbitrap (NCE: 55; resolution: 45,000; normalized AGC target: 200%; maximum injection time: 200 ms; scan range: 100–500 m/z).

#### Data Analysis

Raw data were processed using Proteome Discoverer (v3.0; Thermo Fisher Scientific). MS2 spectra were searched against the SwissProt mouse protein database using the following parameters: MS1 and MS2 mass tolerances were set to 10 ppm and 0.6 Da, respectively. Carbamidomethylation of cysteine (+57.02146 Da) and TMTpro labeling of lysine residues and peptide N-termini (+304.2071 Da) were set as static modifications. Oxidation of methionine (+15.9949 Da) was included as variable modification. Peptide and protein identifications were filtered to a false discovery rate (FDR) of ≤1% at both the peptide and protein levels. Reporter ion quantification was carried out using MS3 spectra meeting the following quality thresholds: average signal-to-noise ratio ≥10 and isolation interference <50%.

The resulting *Baf53b-Cre* dataset included 7581 proteins, and the *Sox10-Cre* dataset included 7012 proteins. Principal Component Analysis was conducted on log2-transformed normalized abundance values for proteins with data in more than 90% of samples (7223 in *Baf53b-Cre* and 6638 in *Sox10-Cre*) using the factoextra package in R [35]. Fold-change and p-value (via two-sample t-test) were calculated for proteins with at least two non-NA/non-zero normalized abundance values. The cutoff for differentially expressed proteins (DEPs) was p < 0.05 and |log2FC| > 0.5849, representing a fold-change cutoff of 1.5. Phenotypic protein changes were significant by p < 0.05 and |log2FC| > 0.5849 in the cKI; Cre- vs. WT; Cre-comparison. Cre-driven protein changes were defined from a list of proteins not significantly altered between cKI; Cre- and WT; Cre- mice by either the p-value or log2FC cutoff that were significantly altered in both cKI; Cre+ vs. cKI; Cre- mice and cKI; Cre+ vs. WT; Cre- mice. Rescue was defined as a protein with a log2FC < -0.152 or log2FC > 0.138 (a 10% change) in cKI; Cre+ mice relative to cKI; Cre- mice. Functional enrichment analysis was conducted with g:Profiler for the genes coding for rescued proteins, with results presented for GO terms [31,32].

### Statistics

Statistical analyses were performed in GraphPad Prism and RStudio. Where applicable, normality was confirmed using the Shapiro-Wilk method and outliers were excluded via ROUT before further testing. Statistical tests and sample size are noted in figure legends. Graphs display the mean ± SEM with the following significance levels: no symbol = not significant, * p < 0.05, ** p < 0.01, *** p < 0.001, or **** p < 0.0001.

## Results

### Genetically silencing mutant ATXN3 in neurons and oligodendrocytes partially attenuates SCA3 motor phenotypes

To define the cell type–specific contributions of mutant ATXN3 to SCA3 phenotypes, we crossed our conditional “disease-off” SCA3 mouse model (cKI) with three Cre drivers: *Baf53b-Cre* to constitutively target neurons (Baf53b), *Sox10-iCre/ERT2* to inducibly target oligodendrocytes and Schwann cells (iSox10), and *Sox10-Cre* to constitutively target both of these populations – neural crest-derived cells and oligodendrocytes (cSox10). Cre- mediated recombination selectively ablates expression of the mutant (Q300) *Atxn3* allele, without affecting expression of the wild type (Q6) *Atxn3* allele. Because *Atxn3*-Knockout mice have no alteration of motor phenotypes [7,36] nor of oligodendrocyte maturation impairments [13,14], phenotypes in Cre+ cKI mice are unlikely to reflect *Atxn3* haploinsufficiency.

Before assessing the contribution of mutant ATXN3 to SCA3 motor phenotypes in each Cre line, we characterized recombination patterns using Ai14 tdTomato reporter mice [24]. Consistent with prior studies, the Baf53b, iSox10 + tamoxifen, and cSox10 drivers produced robust recombination in the expected regions and cell populations of the brain, spinal cord, and dorsal root ganglia (DRG) (**Fig S1-S6**) [21–23,37,38]. In Baf53b mice, recombination was evident in neurons within SCA3-vulnerable regions, including the deep cerebellar nuclei (DCN), Purkinje cells of the cerebellar lobules, and pontine nuclei (**Fig. S2**). In iSox10 mice injected with tamoxifen at 12 weeks, selective recombination in oligodendrocytes of the DCN was observed with no qualitative targeting of neurons, astrocytes, or microglia (**Fig. S4**). Basal Cre activity was observed in vehicle-treated iSox10 mice, but did not localize to oligodendrocytes. In cSox10 mice, recombination was detected in oligodendrocytes and neural crest-derived populations, including Schwann cells and neurons in the DRG, as well as Purkinje cells in the cerebellum (**Fig. S5- S6**). As in tamoxifen-treated iSox10 mice, recombination in astrocytes and microglia was minimal in cSox10 mice (**Fig. S6**).

To confirm Cre-mediated recombination and mutant ATXN3 target engagement, we immunoblotted for ATXN3 in cerebellar lysate from late-stage, 36-week-old mice (**Fig. S7**). As expected, expression of wildtype Q6 ATXN3 was reduced by approximately 50% in cKI mice relative to WT controls in all models. Mutant Q300 ATXN3 was variably reduced from cKI levels in Cre+ mice, with a 60% reduction in Baf53b mice (**Fig. S7c**), no significant reduction in iSox10 + tamoxifen mice (**Fig. S7f**), and a 40% reduction in cSox10 mice (**Fig. S7i**). The lack of a significant change in iSox10 + tamoxifen mice was not surprising, as oligodendrocytes make up a small proportion of total cerebellar cells: approximately 8% when measured by immunohistochemistry [39] or 3% by single-nucleus RNA sequencing [40–42]. Orthogonal validation of target engagement via PCR in the frontal cortex of 36-week-old iSox10 mice confirmed that the mutant ATXN3 allele was indeed edited in cKI mice injected with tamoxifen (**Fig. S8**).

Having established the recombination patterns and mutant ATXN3 target engagement in each model, we next assessed the contribution of the targeted cell populations to SCA3-associated motor deficits. For each model, locomotor activity was measured in the open field and balance and coordination were evaluated on the balance beam at early- (11 or 12 weeks), mid- (24 weeks), and late- (36 weeks) stage disease timepoints (**Fig 1**). In Baf53b mice, neuronal silencing of mutant ATXN3 partially attenuated the reduction of total locomotion at mid-stage disease, but this effect was not sustained at the late-stage timepoint (**Fig. 1b**). Balance beam traversal latency was not modified by neuronal targeting in the Baf53b mice (**Fig. 1c**). In iSox10 administered tamoxifen at 12 weeks, silencing mutant ATXN3 in oligodendrocytes and Schwann cells did not alter motor behavior in the open field or on the balance beam relative to cKI vehicle-treated controls (**Fig. 1d–f**). We observed tamoxifen-associated toxicity in cKI; Cre+ mice, with four of 34 total injected animals dying before 14 weeks of age. No mortality was observed among tamoxifen-treated WT; Cre+ mice or vehicle-injected groups. In contrast, constitutive targeting of neurons, oligodendrocytes, and Schwann cells in cSox10 mice partially rescued open field total locomotion through late-stage disease and fully rescued balance beam traversal latency (**Fig. 1g–i**). These findings suggest that targeting multiple cell populations, and potentially modifying interactions between them, produces a synergistic, sustained rescue of motor phenotypes in SCA3 compared to targeting neurons or myelinating glia alone. Mutant ATXN3 silencing did not modify the body weight deficit in cKI; Cre- mice of the Baf53b or cSox10 lines, and cKI-Veh mice in the iSox10 line did not have reduced weight compared to WT counterparts (**Fig. S9**). Finally, there were no consistent disease- or rescue-associated changes in open field rearing or anxiety-like behavior, or latency to fall on the accelerated rotarod task (**Fig. S9**).

**Fig. 1.**
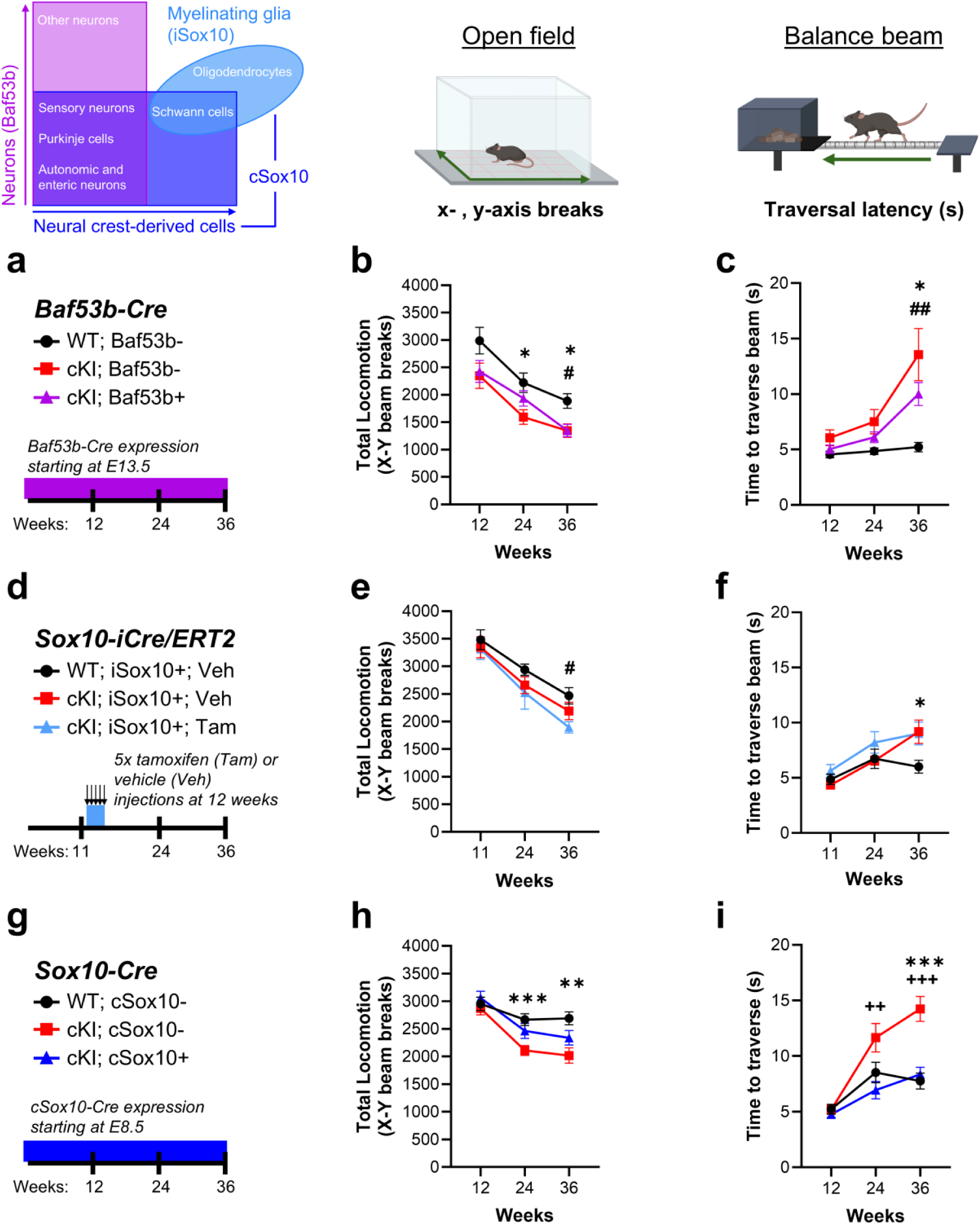
Genetically silencing mutant ATXN3 in neural crest-derived populations and oligodendrocytes produces sustained attenuation of SCA3 motor phenotypes. **(a)** Schematic of behavioral assessment timeline in *Baf53b-Cre* (Baf53b) mice. **(b)** Total locomotion (open field apparatus x-y beam breaks) in WT; Baf53b-, cKI; Baf53b-, and cKI; Baf53b+ mice at 12, 24, and 36 weeks of age. **(c)** Traversal latency (time to traverse the balance beam in seconds) in Baf53b mice at the same timepoints. **(d)** Schematic of behavioral assessment timeline in *Sox10-iCre/ERT2* inducible (iSox10) mice, with Cre induction via tamoxifen (Tam) at 12 weeks. (e) Total locomotion in WT; iSox10+; Vehicle (Veh), cKI; iSox10+; Vehicle, and cKI; iSox10+; Tam mice at 11, 24, and 36 weeks of age. **(f)** Balance beam traversal latency in iSox10 mice at the same timepoints. **(g)** Schematic of behavioral assessment timeline in *Sox10-Cre* constitutive (cSox10) mice, with Cre expression starting at E8.5. **(h)** Total locomotion in WT; cSox10-, cKI; cSox10-, and cKI; cSox10+ mice at 12, 24, and 36 weeks of age. (i) Balance beam traversal latency in cSox10 mice at the same timepoints. *Baf53b-Cre:* n=16– 24 mice/group at 12wk, 14–24 mice/group at 24wk, and 13–18 mice/group at 36wk. *Sox10-iCre/ERT2*: n=9–14 mice/group at 11wk, n=7–15 mice/group at 24wk, and n=7–13 mice/group at 36wk. *Sox10-Cre*: n=24–41 mice/group at 12wk, n=26–34 mice/group at 24wk, and n=13–19 mice/group at 36wk. Graphs display mean ± SEM. Mixed-effects analysis with Tukey’s multiple comparisons test. * indicates the WT; Cre- vs. KI; Cre-comparison for the *Baf53b-Cre* and *cSox10-Cre* models, and WT; iSox10+; Veh vs. cKI; iSox10+; Veh for the *Sox10-iCre/ERT2* model. # indicates the WT; Cre-vs. KI; Cre+ comparison for the *Baf53b-Cre* and *cSox10-Cre* models, and WT; iSox10+; Veh vs. cKI; iSox10+; Tam for the *Sox10-iCre/ERT2* model. + indicates the cKI; Cre+ vs. cKI; Cre-comparison for the *Baf53b-Cre* and *cSox10-Cre* models, and cKI; iSox10+; Tam vs. cKI; iSox10+; Veh comparison for the *Sox10-iCre/ERT2* model. No symbol = not significant; one symbol, p < 0.05; two symbols, p < 0.01; three symbols, p < 0.001.

### Gene and protein expression analyses reveal pathways altered by cell type-specific mutant ATXN3 silencing

Our previous studies correlated the onset of motor phenotypes with the onset of reduced mature oligodendrocyte transcripts in disease-vulnerable brain regions of transgenic SCA3 mice [15]. We also found that broad reduction of brain ATXN3 expression via anti-*ATXN3* antisense oligonucleotides ameliorates motor phenotypes and oligodendrocyte maturation abnormalities [12,16]. These findings raised the possibility that the sustained motor rescue observed in cSox10 mice resulted from restoration of oligodendrocyte maturation. To test this hypothesis, we first examined mature oligodendrocyte transcripts from the cerebellum of cSox10 mice at early- and mid-disease stages (**Fig. 2a-b**). In agreement with prior work from our lab, the mature oligodendrocyte transcripts *Plp1*, *Mobp*, *Mal,* and *Ugt8a* were significantly reduced in cKI; Cre- mice relative to WT controls at both timepoints [15]. Unexpectedly, these transcripts remained reduced in cKI; cSox10+ mice, indicating that silencing mutant ATXN3 in oligodendrocytes and neural crest-derived cells did not alter this transcriptional signature of oligodendrocyte maturation. Thus, the sustained motor rescue in cSox10 mice occurred independently of mature oligodendrocyte transcriptional normalization. Moreover, the persistent reduction of mature oligodendrocyte transcripts despite oligodendrocyte targeting suggests that non-cell-autonomous mechanisms likely contribute to oligodendrocyte maturation abnormalities in SCA3.

**Fig. 2.**
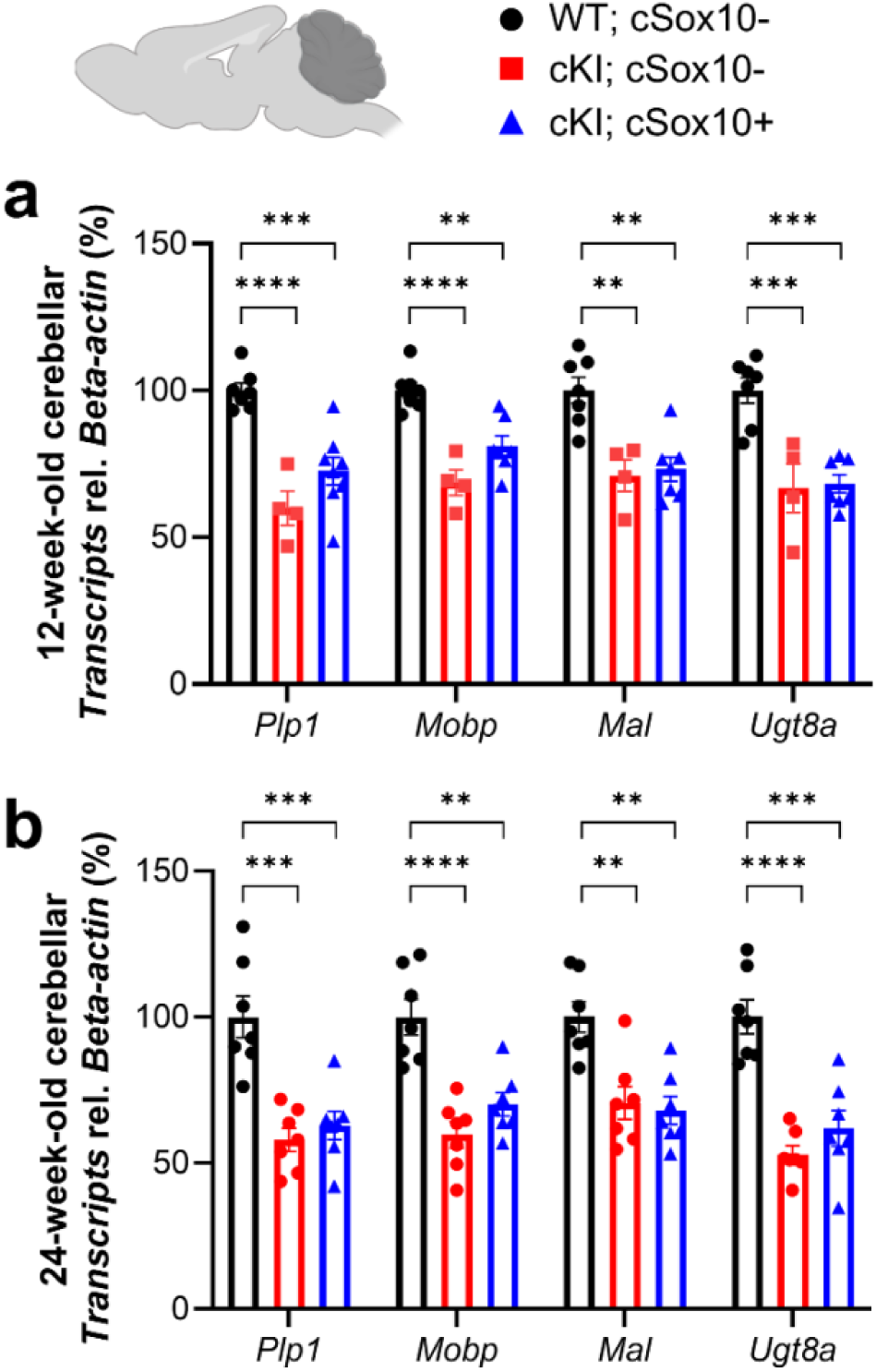
Mature oligodendrocyte transcripts are not rescued by silencing mutant ATXN3 in neural crest-derived cells and oligodendrocytes at early- or mid-stage SCA3 disease. Quantification of the mature oligodendrocyte transcripts *Plp1, Mobp, Mal, and Ugt8a* in the cerebellum of WT; cSox10-, cKI; cSox10-, and cKI; cSox10+ mice at **(a)** 12 weeks and **(b)** 24 weeks of age. Each transcript is normalized to *Beta-actin* and to the average of WT; cSox10- mice, with n=4–8 mice/group at 12 weeks and n=7 mice/group at 24 weeks. Graph represents mean ± SEM, with each point as one mouse. One-way ANOVA with post- hoc Tukey’s multiple comparisons test. No symbol = not significant; ** p < 0.01; *** p < 0.001; **** p < 0.0001.

We therefore broadened our search for molecular correlates of motor rescue by performing unbiased bulk RNA sequencing and proteomic profiling of the cerebellum from 36-week-old Baf53b and cSox10 mice (**Fig. 3, S10**). Principal component analysis (PCA) of the 500 most differentially expressed genes in WT; Cre-, cKI; Cre-, and cKI; Cre+ mice of the Baf53b and cSox10 lines revealed minimal separation of samples by genotype (**Fig. S10a, b**). In the Baf53b cohort, 131 genes were differentially expressed in cKI; Baf53b- mice compared to WT controls, of which 109 (83%) rescued, meaning shifted toward WT levels following neuronal silencing of mutant ATXN3 (**Fig. 3b, S10c**). Rescue was defined as a change of at least 10% in log2 fold change (log2FC) of a gene in cKI; Cre+ mice relative to cKI; Cre- mice, in the direction of WT expression. In the cSox10 cohort, 47 genes were differentially expressed in cKI; cSox10-mice relative to WT controls, of which 34 (72%) shifted toward WT levels following mutant ATXN3 silencing in cKI; cSox10+ mice (**Fig. 3c, S10d**). Gene ontology enrichment analysis associated rescued genes in the Baf53b mice with the plasma membrane, cell surface receptor signaling pathways, and the cell periphery (**Fig 3d**). In cSox10 mice, the top enriched terms were: response to other organism, response to external biotic stimulus, and response to external stimulus (**Fig 3e**).

**Fig. 3.**
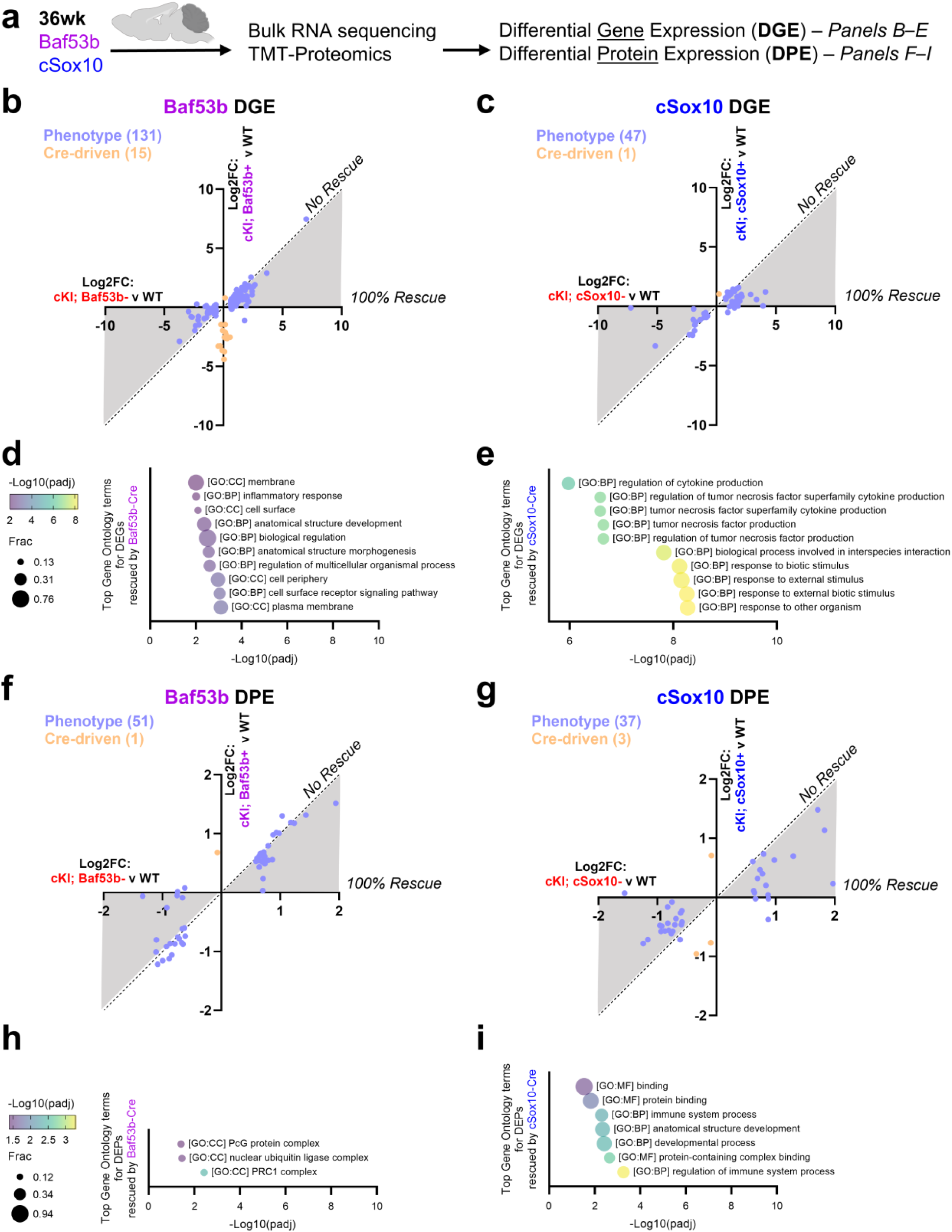
Differential gene and protein expression analyses on the late-stage cerebellum reveal pathways altered by cell type-specific mutant ATXN3 silencing. **(a)** Schematic of experimental design. The cerebellum was isolated from late-stage, 36-week-old *Baf53b-Cre* (Baf53b) and *Sox10-Cre* (cSox10) mice for bulk RNA sequencing and Tandem Mass tag (TMT)-Proteomics to identify differentially expressed genes and proteins, respectively. Subsequent plots in **b**, **c**, **f**, and **g** compare the log2 fold-change (FC) in cKI; Cre- mice relative to WT mice on the x-axis and the log2FC in cKI; Cre+ mice relative to WT mice on the y-axis. Phenotype represents significantly altered genes in cKI; Cre- vs. WT mice. Cre-driven represents significantly altered genes in the cKI; Cre+ vs. cKI; Cre- and cKI; Cre+ vs. WT two-way comparisons, that do not meet differential expression thresholds in the cKI; Cre- vs. WT two-way comparison. The shaded area represents rescued genes, with percent rescue in cKI; Cre+ mice in the direction of WT decreasing as a point approaches the dotted line. **(b)** Differential gene expression analysis in Baf53b mice. **(c)** Differential gene expression analysis in the cSox10 mice. **(d)** The top ten statistically significant Gene Ontology (GO) terms derived from the genes rescued by Baf53b. Frac represents the fraction of genes in the GO term contained in the dataset. **(e)** The top ten GO terms from the genes rescued by cSox10. **(f)** Differential protein expression analysis in the *Baf53b- Cre* model. **(g)** Differential protein expression analysis in the *cSox10-Cre* model. **(h)** The three significant GO terms derived from the genes coding for proteins rescued by Baf53b. **(i)** The seven significant GO terms derived from the genes coding for proteins rescued by cSox10. n=3–4 mice/group for RNAseq and n=4–6 mice/group for proteomics. The cutoff for differential gene expression was p-adj < 0.05 and |log2FC| > 0.5849. The cutoff for differential protein expression was p < 0.05 and |log2FC| > 0.5849. Rescue was defined as a gene or protein with a log2FC < - 0.152 or log2FC > 0.138 (a 10% change) in cKI; Cre+ mice relative to cKI; Cre- mice, in the direction of WT; Cre-. BP = Biological Process; CC = Cellular Component; MF = Molecular Function.

We next examined whether cell type–specific mutant ATXN3 silencing produced corresponding changes at the protein level. PCA of the proteomics data similarly showed no clear genotype-dependent separation in Baf53b mice (**Fig. S10e**). In contrast, cSox10 mice showed moderate separation by genotype, with cKI; Cre+ mice displaying an intermediate proteomic profile between WT and cKI; Cre- controls (**Fig. S10f**). Differential expression analysis identified 51 proteins significantly altered in cKI; Baf53b-mice relative to WT (**Fig. 3f**) and 37 altered proteins in cKI; cSox10-relative to WT (**Fig. 3g**). Although the numbers of disease-associated proteins were comparable, their extent of rescue differed markedly: 31% (16/51) shifted toward WT in Baf53b mice, compared with 89% (33/37) in cSox10 (**Fig. S10g-h**). Functional enrichment analysis on the genes encoding the rescued proteins revealed only three terms associated with rescue in Baf53b mice: PRC1 complex, nuclear ubiquitin ligase complex, and PcG protein complex, suggesting changes in chromatin and protein quality control (**Fig. 3h**). In cSox10 mice, the top enriched terms included regulation of immune system process, protein-containing complex binding, and developmental process (**Fig. 3i**).

Together, transcriptomic and proteomic analyses highlight that mutant ATXN3 silencing in oligodendrocytes and neural crest-derived cells partially normalize immune-related molecular signatures in the cerebellum. Notably, this rescue occurred despite minimal recombination in astrocytes and microglia of the cSox10 mice (**Fig S6**). This prompted us to interrogate protein-level changes across cSox10-targeted (neurons, oligodendrocytes) and non-targeted (astrocytes, microglia) cell populations in the cerebellum.

### Silencing mutant ATXN3 in cSox10-targeted populations modifies astrocytic and microglial protein signatures

To elucidate the cellular context of these immune-related changes, we queried the proteomics data from 36-week-old Baf53b and cSox10 mice for proteins enriched in neurons, oligodendrocytes, astrocytes, and microglia (**Fig 4**). Literature-supported neuron-enriched proteins, including the pan-neuronal markers RBFOX3 and MAP2 and the Purkinje cell-associated proteins CALB1, PCP2, PVALB, and ALDOC, were unchanged between cKI; Cre- mice and WT controls in both models (**Fig. 4a**) [43–49]. Mature oligodendrocyte markers were reduced in cKI; Cre- mice relative to WT counterparts of both models (**Fig. 4b**), though the only proteins that qualified as differentially expressed with the cutoff of p < 0.05 and |log2FC| > 0.5849 (representing a FC of 1.5) were ASPA and SERPINB1A. Neither ASPA nor SERPINB1A were rescued in cKI; Baf53b+ mice, but both were partially rescued in cKI; cSox10+ mice. Interestingly, SERPINB1A, which is enriched in myelinating oligodendrocytes, was recently reported to be reduced in a Knock-in mouse model of SCA2, another polyglutamine SCA [50]. The failure of either Cre model to rescue PLP1, MOBP, or UGT8A in the cerebellum of late-stage cKI mice is consistent with the qPCR results of mature oligodendrocyte transcriptional dysregulation at early- and mid-stage disease (**Fig. 2**). Additionally, the convergence of transcript and protein data strongly suggests that improvement of motor symptoms in cKI; cSox10+ mice is not driven by altered neuronal or mature oligodendrocyte signatures.

**Fig 4.**
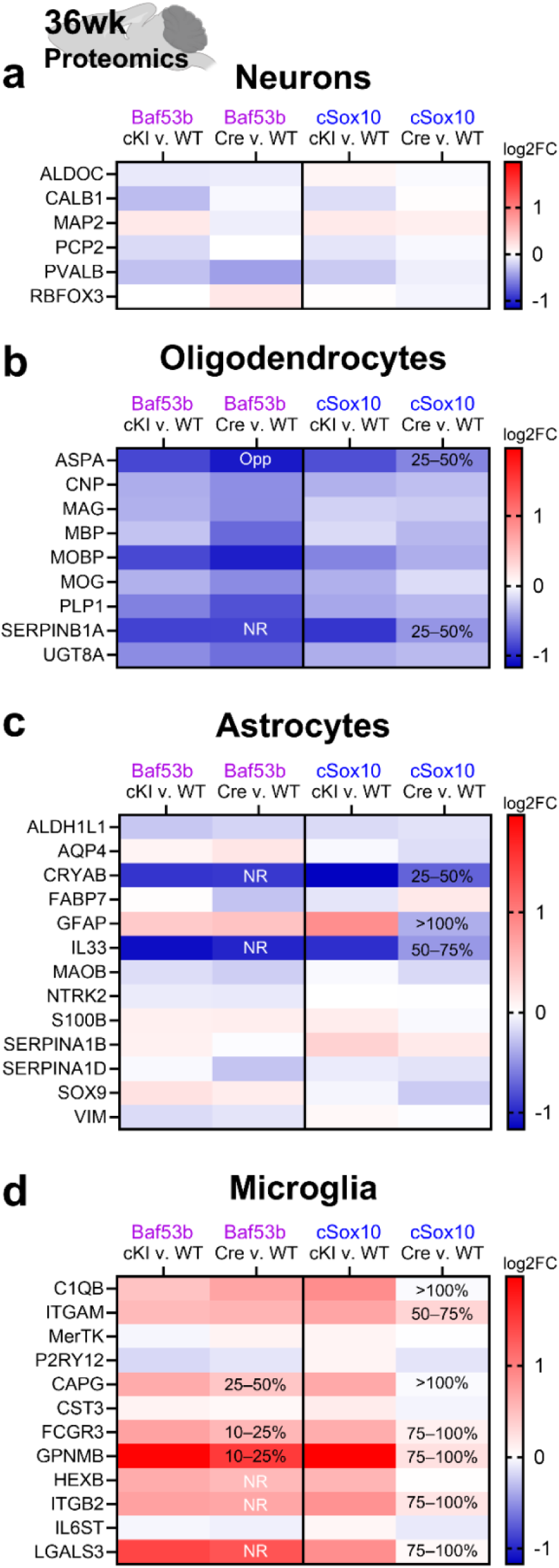
Genetic silencing of mutant ATXN3 has non-cell-autonomous effects on astrocyte and microglia protein markers. Cell type-enriched protein markers were profiled in the 36-week cerebellar dataset from Baf53b and cSox10 mice. Each heatmap displays the log2FC in cKI; Cre- versus WT; Cre- mice (cKI v. WT) and cKI; Cre+ versus WT; Cre- mice (Cre v. WT). Baf53b is on the left, and cSox10 is on the right. If a protein was differentially expressed in cKI; Cre- mice relative to WT controls (p < 0.05 and |log2FC| > 0.5849), the rescue category is displayed on the heatmap. Rescue was defined as a protein with a minimum 10% change in cKI; Cre+ mice relative to cKI; Cre- mice, in the direction of WT; Cre-. NR indicates not rescued proteins, and Opp indicates those changed in the opposite direction of WT. Heatmaps display proteins enriched in **(a)** neurons, **(b)** oligodendrocytes, **(c)** astrocytes, and **(d)** microglia. n=4–6 mice/group.

In contrast to the consistent lack of change in neuron- and oligodendrocyte-enriched proteins across Baf53b and cSox10 mice, profiling astrocyte- and microglia-enriched proteins highlighted a diverging pattern of rescue between the two Cre models (**Fig 4c-d**). Although the homeostatic astrocyte markers ALDH1L1 and AQP4 were unchanged between genotypes in both models, the reactivity-associated proteins CRYAB and IL33 were significantly reduced in cKI; Cre- mice relative to WT counterparts (**Fig. 4c**) [51,52]. These proteins were partially rescued in cKI; cSox10+ mice, but not in cKI; Baf53b+ mice. Similarly, GFAP was also increased in cKI; cSox10- mice and restored to WT levels only in cKI; cSox10+ mice. Microglia-enriched proteins CAPG, FCGR3, GPNMB, ITGB2, and LGALS3 were increased in cKI; Cre- mice of both models relative to WT controls (**Fig. 4d**) [53–56]. Among these five proteins, three (CAPG, FCGR3, and GPNMB) were partially rescued by Baf53b-Cre, while all were partially or fully rescued by cSox10-Cre. Together, these findings indicate that cSox10-Cre more broadly normalizes overt astrocyte- and microglia-associated inflammatory signatures, likely through non-cell-autonomous mechanisms.

### Cerebellar alternative splicing dysregulation is partially rescued by cell type-specific mutant ATXN3 silencing

Because differential gene and protein expression revealed relatively modest rescue, particularly in the directly targeted cell populations, we next examined alternative RNA splicing. Prior studies identified splicing as a sensitive measure of disease-associated molecular changes and rescue in SCA3-vulnerable tissues [7,17,18]. Using rMATS, we quantified dysregulation across skipped exon (SE), retained intron (RI), mutually exclusive exon (MXE), alternative 3’ splice site (A3SS) and alternative 5’ splice site (A5SS) events. Across these event classes, we identified 1832 dysregulated events (ΔPSI>10%, FDR <0.05) in cKI; Baf53b-relative to WT; Baf53b-mice, of which 1244 were rescued by cKI; Baf53b+ (**Fig. 5a**). Rescue was defined as a PSI change in the cKI; Cre+ mice, >10% from cKI; Cre- mice, in the direction of WT; Cre- mice, with an absolute value of at least 5%. In cSox10 mice, 1680 events were dysregulated in cKI; cSox10-relative to WT; cSox10-mice, of which 965 were rescued in cKI; cSox10+ mice (**Fig. 5b**).

**Fig. 5.**
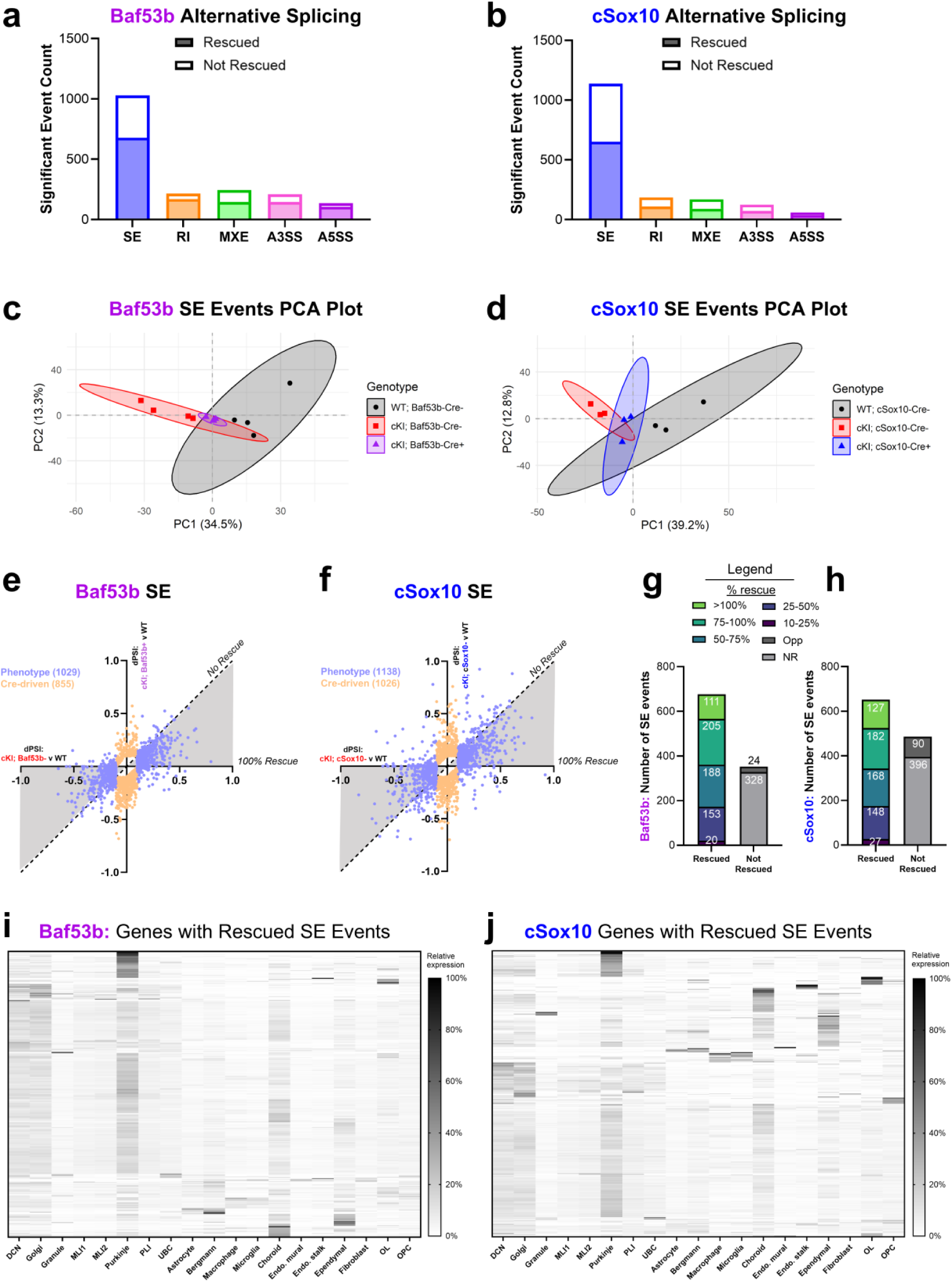
Alternative splicing dysregulation in the late-stage cerebellum is partially attenuated by mutant ATXN3 cell type-specific silencing. Number of significant mis-spliced events (including rescued and not rescued) in the cerebellum of 36-week-old cKI; Cre- vs. WT mice in the categories of skipped exon (SE), retained intron (RI), mutually exclusive exons (MXE), alternative 3’ splice site (A3SS), and alternative 5’ splice site (A5SS) for the (a) *Baf53b-Cre* (Baf53b) and (b) *Sox10-Cre* (cSox10) models. **(c)** Principal component analysis (PCA) plot based on SE events in Baf53b mice, where each point is one mouse and a 95% confidence ellipse surrounds each genotype. **(d)** PCA plot based on SE events in cSox10 mice. **(e)** Plot for Baf53b SE events displaying the delta percent spliced in (dPSI) in cKI; Cre- relative to WT mice on the x-axis, and dPSI in cKI; Cre+ mice vs. WT mice on the y-axis. Phenotype represents significant SE events in cKI; Cre- vs. WT mice. Cre-driven represents significant SE events in the cKI; Cre+ vs. cKI; Cre- and cKI; Cre+ vs. WT two-way comparisons, that do not meet differential expression thresholds in the cKI; Cre- vs. WT two-way comparison. The shaded area represents rescued events, with percent rescue in cKI; Cre+ mice in the direction of WT decreasing as a point approaches the dotted line. The cutoff for SE event significance was FDR < 0.05 and dPSI > 10%. **(f)** Plot for cSox10 SE events displaying dPSI in cKI; Cre- vs. WT on the x-axis and cKI; Cre+ vs. WT on the y-axis. **(g)** Count of rescued and not rescued SE events in cKI; Baf53b+ mice, with percent rescue toward WT color-coded. **(h)** Count of rescued and not rescued SE events in cKI; cSox10+ mice. **(i)** Relative expression of significant events across cerebellar cell types in the Baf53b dataset. Color scale represents the relative expression of the SE event gene in each cell type. **(j)** Relative expression of significant events across cerebellar cell types in the cSox10 dataset. n=3–4 mice/group for RNAseq. Rescue was defined as an average PSI in cKI; Cre+ mice at least 10% different from cKI; Cre- mice, in the direction of WT; Cre- controls. DCN = deep cerebellar nuclei, MLI1 = molecular layer interneuron 1, MLI2 = molecular layer interneuron 2, PLI = Purkinje layer interneuron, UBC = unipolar brush cell, Endo. = endothelial, OL = oligodendrocyte, OPC = oligodendrocyte precursor cell.

Consistent with previous work in the SCAs [7,17,18,57,58], SE events constitute the majority of dysregulated and rescued events in each model. We therefore proceeded with deeper analysis of SE events. PCA plots based on SE events in Baf53b and cSox10 mice showed more distinct clustering by genotype than the plots for differential gene or protein expression. We observed strong separation between WT; Cre- and cKI; Cre- mice of both lines, with cKI; Cre+ groups clustering between controls (**Fig. 5c-d**), consistent with partial rescue of SE dysregulation. There were 1029 significant SE events in cKI; Cre- mice relative to WT; Cre- mice of the Baf53b model (**Fig. 5e**) and 1138 in the cSox10 model (**Fig. 5f**). Of these, 677 events (66%) were rescued in cKI; Baf53b+ cerebella (**Fig. 5g**) and 652 events (57%) were rescued in cKI; Sox10+ cerebella (**Fig. 5h**). There was no bias toward exon inclusion or exclusion in the significant or rescued events (**Fig. S11a-b**).

To understand the cellular context of the rescued events, we performed cell type-enrichment analysis of SE-affected genes using cerebellar data from the Allen Brain Cell Atlas [59]. Consistent with the recombination patterns of the Cre drivers, genes containing rescued SE events rescued in Baf53b mice were enriched in neuronal populations, including Purkinje cells and DCN (**Fig 5i**). Genes containing SE events rescued by cSox10 were enriched in these neuronal populations as well as oligodendrocytes (**Fig. 5j**). Because the reference atlas used for this analysis comes from healthy mouse brains, disease-associated astrocytic and microglial signatures may be underrepresented in this analysis. Gene ontology enrichment analysis revealed that the top terms associated with SE event genes in Baf53b and cSox10 mice involve cytoskeleton and ion channel function (**Fig. S11c-d**). Together, these findings highlight RNA splicing as a putative modulator of neuron and oligodendrocyte function in SCA3, and support cell autonomous mutant ATXN3 lowering strategies as a viable way to ameliorate dysfunctional splicing.

## Discussion

SCA3 affects multiple neuronal and glial populations, but the extent to which mutant ATXN3 expression within specific cell types drives functional disease impairments remained unclear. This study harnessed a novel conditional mouse to silence mutant ATXN3 in three populations: neurons, oligodendrocytes and Schwann cells, or a broader combination of oligodendrocytes and neural crest–derived populations. We found the greatest magnitude of behavioral rescue following mutant ATXN3 silencing in neural crest-derived cells and oligodendrocytes under the *Sox10-Cre* driver, with a partial rescue of total locomotion and a full rescue of balance beam traversal latency.

Cerebellar transcriptomic and proteomic analysis showed that this behavioral attenuation occurred without restoration of mature oligodendrocyte signatures, but was associated with partial normalization of overt astrocyte- and microglia-related inflammatory signals. Cell type-specific mutant ATXN3 silencing also rescued numerous disease-associated RNA-splicing abnormalities, with rescued events enriched in cell populations targeted by each Cre driver. Overall, these findings reveal both cell-autonomous and non-cell-autonomous contributions to SCA3 and suggest that effective therapies may require broad cellular delivery, with neuroinflammatory and RNA-splicing signatures as candidate molecular correlates of therapeutic response.

The differential behavior rescue across the three Cre models highlights motor phenotypes arising from mutant ATXN3 in specific cell populations. The greater magnitude and durability of rescue in cSox10 mice may reflect the combined effects of reducing mutant ATXN3 across interacting central and peripheral cell populations, as we recently reported phenotypic rescue resulting from mutant ATXN3 silencing in sensory neurons of the peripheral nervous system [7]. These findings extend previous work showing that broad CNS reduction of ATXN3 with anti-*ATXN3* antisense oligonucleotides improves motor function in SCA3 transgenic mice [12,16] and support the development of therapeutic strategies capable of reaching multiple disease-relevant cell types.

Unexpectedly, silencing mutant ATXN3 in neural crest-derived populations and oligodendrocytes did not restore mature oligodendrocyte transcript or protein signatures, suggesting that oligodendrocyte maturation deficits are not exclusively cell autonomous. Our previous evidence for a cell-autonomous maturation deficit was based on primary oligodendrocyte precursor cells derived from one-week-old YACQ84 transgenic SCA3 mice [13,14]. The present *in vivo* findings suggest a more complex model in which cell-intrinsic mutant ATXN3 interacts with signals from the surrounding tissue environment, potentially including the persistent inflammatory signatures observed here (**Fig. 4**), to impair oligodendrocyte maturation. Conversely, the stress of *ex vivo* primary cell culture may accentuate mutant ATXN3 toxicity and oligodendrocyte maturation deficits. This possibility is consistent with the absence of detectable oligodendrocyte maturation abnormalities *in vivo* at 3 weeks of age and their emergence by 4 weeks in YACQ84 mice [15]. Distinguishing between these mechanisms will require directly testing whether mutant ATXN3 expression in oligodendrocytes is sufficient to impair their maturation *in vivo*. Such studies will require development of a conditional “ON” SCA3 KI mouse model. Using the currently available tools, future studies could cross cKI mice with astrocyte- or microglia-specific Cre drivers to selectively silence mutant ATXN3 in each cell type and survey changes to oligodendrocyte maturation, which would clarify the role of these cells in the oligodendrocyte maturation impairment.

Our findings add to the growing body of evidence distinguishing RNA-splicing dysregulation as a prominent molecular feature of SCAs [7,17,18,57,58]. Genes containing splicing events rescued by Baf53b mice were enriched in the cerebellar neuronal populations, whereas those rescued by cSox10 mice were enriched in both neurons and oligodendrocytes. This supports a cell-autonomous contribution of mutant ATXN3 to splicing dysfunction. Moreover, the extensive rescue of alternative splicing abnormalities, despite comparatively modest changes in gene and protein abundance, suggests that RNA splicing may be a sensitive molecular readout of mutant ATXN3 target engagement and therapeutic response. Nevertheless, the functional consequences of these events and their relationship to behavioral rescue remain to be determined.

Several limitations should guide the interpretation of these findings and future studies. First, while the conditional mouse model avoids the confounds of transgene overexpression, it expresses and silences murine *Atxn3* rather than the human gene, potentially limiting direct translation to patients with SCA3. Second, the timing and durability of mutant ATXN3 silencing in iSox10 mice may have limited our ability to detect behavioral rescue if tamoxifen administration at 12 weeks of age occurred after relevant pathogenic processes were established. In addition, turnover of targeted cells could have introduced newly generated cells that continued to express mutant ATXN3, reducing the durability of target engagement. Although our induction timepoint was selected to succeed the restriction of *Sox10* expression to myelinating glia while still preceding severe disease manifestations, future studies using earlier or repeated tamoxifen dosing, or an alternative oligodendrocyte-selective Cre driver, could clarify the questions surrounding timing and sustained silencing. Finally, the limited correspondence between cerebellar differential gene or protein expression and behavioral rescue suggests that analysis of additional disease-vulnerable regions may be necessary to fully dissect the cell types and molecular processes driving behavioral phenotypes. Examples include the brainstem and spinal cord, which demonstrate robust SCA3 pathogenic signatures [14,17]. Despite these limitations, this study advances our understanding of cell type-specific contributions to SCA3 disease and highlights promising avenues for continued study.

## Summary

In summary, our findings demonstrate that SCA3 phenotypes arise from the coordinated contribution of multiple neuronal and glial populations. Silencing mutant ATXN3 in neurons or myelinating glia alone did not produce sustained motor rescue, whereas broader targeting of oligodendrocytes and neural crest-derived populations yielded more robust and durable behavioral improvement. This rescue occurred without restoration of oligodendrocyte maturation signatures. Instead, behavioral attenuation was associated with non-cell-autonomous normalization of astrocyte- and microglia-related inflammatory signatures and cell-autonomous rescue of RNA-splicing abnormalities. Together, these results underscore the importance of interactions among disease-relevant cell populations and support therapeutic strategies that broadly reduce mutant ATXN3 across the central and peripheral nervous systems.

## Acknowledgements

The authors thank Dr. Andrew Lieberman and Emily Januck for sharing the *Sox10-iCre/ERT2* mouse model. We thank Robert Merritt at the RNA Institute Next Generation Transcriptomic Facility as well as the Center for Functional Genomics, both at the University at Albany, for RNA sequencing expertise and instrumentation. Finally, we thank the Proteomics Resource Facility at the University of Michigan for performing proteomics and for providing the text for the methods describing this experiment. Figure schematics were created using BioRender.

## Declarations

### Material and/or Code availability

Proteomics data are available at MassIVE MSV000103325. Bulk RNA sequencing data of WT; *Baf53b-* and cKI; *Baf53b-* mice is available at GSE347811. Bulk RNA sequencing data of all other mice (cKI; *Baf53b+* mice and all cSox10 mice) are available at GSE348508. All other data are available upon reasonable request of the corresponding author.

### Author Contributions

Alexandra F. Putka, Jacen Emerson, and Hayley S. McLoughlin contributed to the study conception and design. Material preparation, data collection, and analysis were performed by all authors. The first draft of the manuscript was written by Alexandra F. Putka, Jacen Emerson, and Hayley S. McLoughlin. The final manuscript was read and approved by all authors.

