## Supplemental Methods and Figures for "Investigating the necessity of neuronal and oligodendroglial mutant ATXN3 expression for Spinocerebellar ataxia type 3 phenotypes"

### Supplementary Material

### Supplementary Methods

#### Immunofluorescence

Following overnight fixation with 4% PFA, tdTomato-positive 36-week *Baf53b-Cre*, 12-week *Sox10-Cre*, and 14-week *Sox10-iCreERT2* brains, spinal cords, and DRG were cryopreserved in 30% sucrose. Brains were sagittally sliced at 25 μm on a Leica SM 200R sliding microtome with a freezing stage and placed free-floating in cryoprotectant solution as previously described [1]. Spinal cords and DRG were sectioned directly onto Superfrost Plus slides (Fisher Scientific, 12-550-15) using a Lecia CM 1850 cryostat at 20 μm and 10 μm, respectively. Staining of all tissues followed the same general procedure as previously described [2]. Sections were briefly washed with PBS, followed by one-hour incubation with M.O.M. Mouse IgG Blocking Reagent (Vector Laboratories, BMK-2202) diluted in 5% normal goat serum in PBS. Sections were PBS-washed again and then incubated in M.O.M. Protein Concentrate in PBS. Primary antibodies (listed in **Supplementary Table 6**) were diluted in the M.O.M. Protein Concentrate buffer and sections were incubated overnight at 4°C. The following day, sections were washed in PBS before a one-hour incubation with secondary antibodies (listed in **Supplementary Table 6**). Finally, sections were stained with DAPI (Sigma-Aldrich, D9564) in PBS and washed. Brain sections were mounted onto slides in PBS, and all tissues were coverslipped with ProLong Gold Antifade Reagent (Invitrogen, P36930). This procedure was optimized for each tissue and mouse strain. All incubation steps for *Baf53b-Cre* mice occurred at 4°C. Brain and spinal cord tissue from *Sox10-Cre*, *Sox10-iCreERT2* mice, and DRG tissue from all three strains, underwent antigen retrieval in 0.01 M sodium citrate buffer (pH 8.5) at 80°C for 30 minutes before washing and blocking. Finally, for Sox10 immunodetection in the brain and spinal cord, 0.3% Triton X-100 (Fisher Scientific, BP151) was included in the blocking solution. Slides were imaged on a Leica Stellaris 5 confocal system, using 10X air, 20X air, and 100X oil objectives, with single-plane or z-stack imaging noted in figure legends. ImageJ was used to combine channels of interest for the figures.

#### Western blot

36-week cerebellar protein samples from *Baf53b-Cre*, *Sox10-Cre*, and *Sox10-iCre/ERT2* mice (20–30 μg) were run on 4-20% SDS-PAGE gels and transferred to 0.45µm nitrocellulose membranes. Total protein load was assessed using Direct Blue (Sigma-Aldrich, 212407) before membranes were blocked and incubated overnight in primary antibody: mouse anti-ATXN3 (1H9, 1:1000, Millipore, MAB5360) or mouse anti-1C2 (1:1000, Millipore, MAB1574). The following day, membranes were washed prior to incubation with peroxidase-conjugated anti-mouse secondary antibody (1:5000, Jackson ImmunoResearch Laboratories) and visualization with SuperSignal West Femto imaging reagents (Thermo Fisher Scientific 34094) using the Syngene G:BOX Mini imaging system. Band intensities were quantified in ImageJ and normalized to Direct Blue intensity between 20 and 75 kDa.

#### *Atxn3* Exon 4 PCR

Genomic DNA was isolated from the frontal cortex of 36-week-old *Sox10-iCre/ERT2* mice using the DNeasy Blood & Tissue Kit (Qiagen 69506). PCR for *Atxn3* exon 4 was conducted with forward primer 5’-CACAGGCACTCAGACACACT-3’ and reverse primer 5’-AGTGATGCTAAGCACAAAGCC-3’. PCR products were resolved via gel electrophoresis, with gel images obtained on a BIORAD GelDoc Go Imaging System.

#### Supplemental mouse behavior assessments

##### Open field apparatus: rearing and anxiety-like behavior

At 12, 24, and 36 weeks of age in the *Baf53b-Cre* and *Sox10-Cre* lines, and 11, 24, and 36 weeks of age in the *Sox10-iCre/ERT2* line, mice were placed in the open field apparatus for 30 minutes (San Diego Instruments). Z-axis beam breaks were recorded as a measure of rearing, and the percentage of the percentage of total locomotion (x-y beam breaks) in the center of the apparatus was recorded as a readout of anxiety-like behavior.

#### Accelerated rotarod

To assay motor coordination, learning, and strength, accelerated rotarod testing was conducted at 12, 24, and 36 weeks in *Baf53b-Cre* and *Sox10-Cre* mice. Mice were tested on a rotarod apparatus (Med Associates Inc.) that accelerated from four to 40 rotations per minute over a five-minute period, with three trails per day (five minutes rest in between) across four consecutive days. The latency for a mouse to fall from the rod on each trial was recorded, and data are shown for the fourth day of testing. As with other behavioral assessments, data from cKI; *Cre+* mice of each strain were compared to sex- and age-matched WT; *Cre-* and cKI; *Cre-* mice.

### Supplementary Tables

**Supplementary Table 1.** Sample information for all mice used in RNAseq and proteomics studies.

**Supplementary Table 2.** Gene expression pairwise comparison data for WT, cKI; Cre-, and cKI; Cre+ mice from cSox10 and Baf53b strains.

**Supplementary Table 3.** Protein expression pairwise comparison data for WT, cKI; Cre-, and cKI; Cre+ mice from cSox10 and Baf53b strains.

**Supplementary Table 4.** Splicing event rescue analysis for each event category in cSox10 and Baf53b strains. avg column represents average PSI in the associated genotype.

**Supplementary Table 5.** Gene ontology results for rescued genes, proteins, and splicing events in Baf53b and cSox10 mice.

**Supplementary Table 6.** Primary and secondary antibody information.

| **Primary Antibody** | **Manufacturer, Catalog #** | **Concentration** | **Secondary antibody**  **(Invitrogen Alexa Fluor at 1:1000)** |
| --- | --- | --- | --- |
| Rat anti-NeuN | Abcam, ab279297 | 1:500 for *Baf53b-Cre* brain/spinal cord  1:200 for all other tissue | 647 goat anti-rat (A21247) |
| Mouse anti-tdTomato | OriGene, TA180009 | 1:500 | 568 goat anti-mouse (A11004) |
| Mouse anti-GFAP | Cell Signaling Technology, 3670 | 1:500 | 488 goat anti-mouse (A11001) |
| Rabbit anti-IBA1 | Invitrogen, PA521274 | 1:200 | 647 goat anti-rabbit (A21244) |
| Rabbit anti-Sox10 | Cell Signaling Technology, 89356 | 1:250 | 488 goat anti-rabbit (A11008) |
| Rabbit anti-S100 beta | Abcam, ab52642 | 1:200 | 488 goat anti-rabbit (A11008) |

### Supplemental Figures

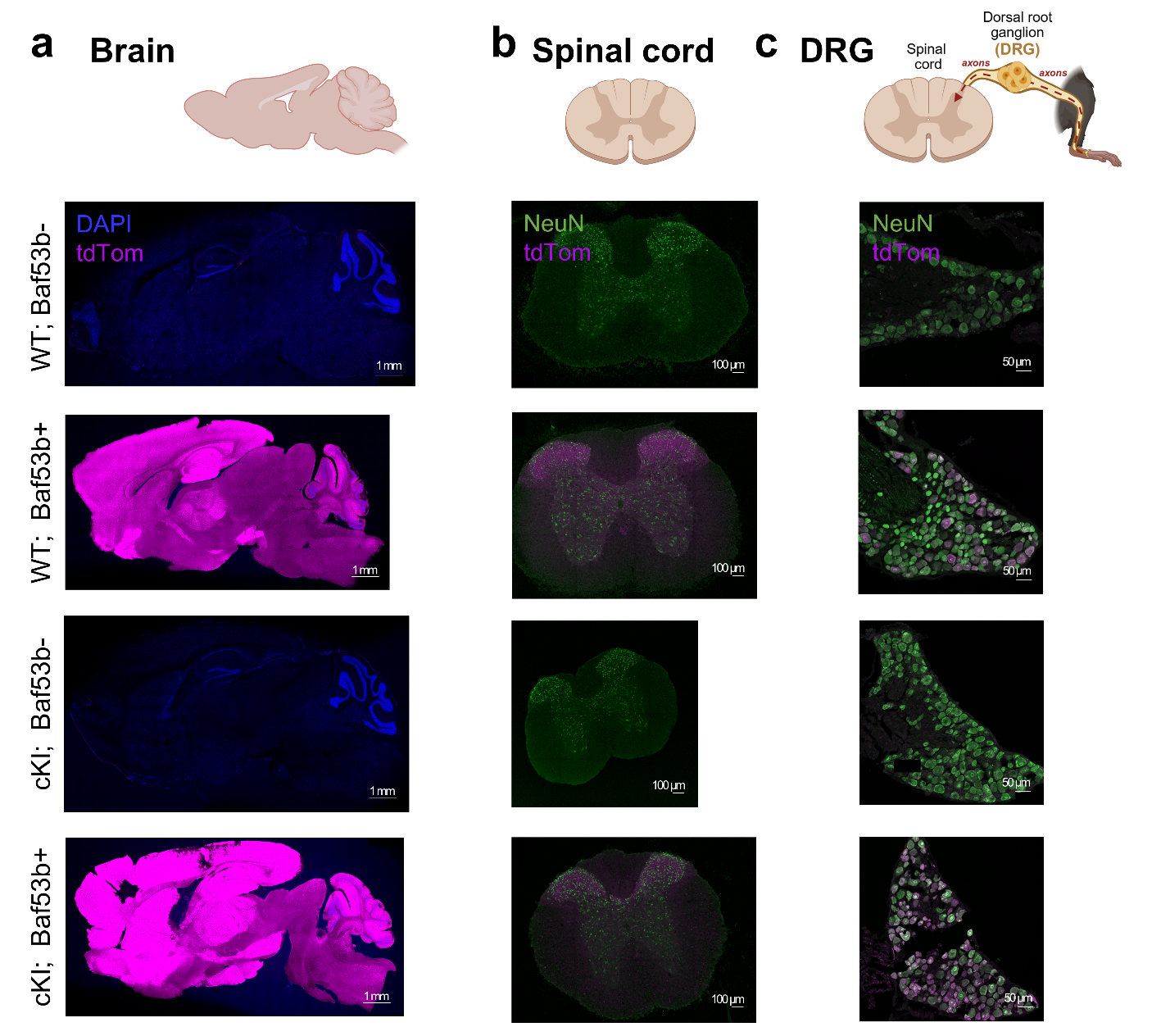
**Fig. S1 *Baf53b-Cre; tdTomato* reporter mice validate robust Cre recombination in the central and peripheral nervous systems**

**(a)** Representative immunofluorescent images of 36-week-old WT; Baf53b-, WT; Baf53b+, cKI; Baf53b-, and cKI; Baf53b+ mice stained for tdTomato (tdTom) and DAPI in sagittal brain sections (scale bar = 1 mm). Representative images of tdTom and NeuN staining in Baf53b **(b)** spinal cord sections (scale bar = 100 µm) and **(c)** DRG sections (scale bar = 50 µm). Sagittal slice images are single-plane, and all others are maximum z-projections.

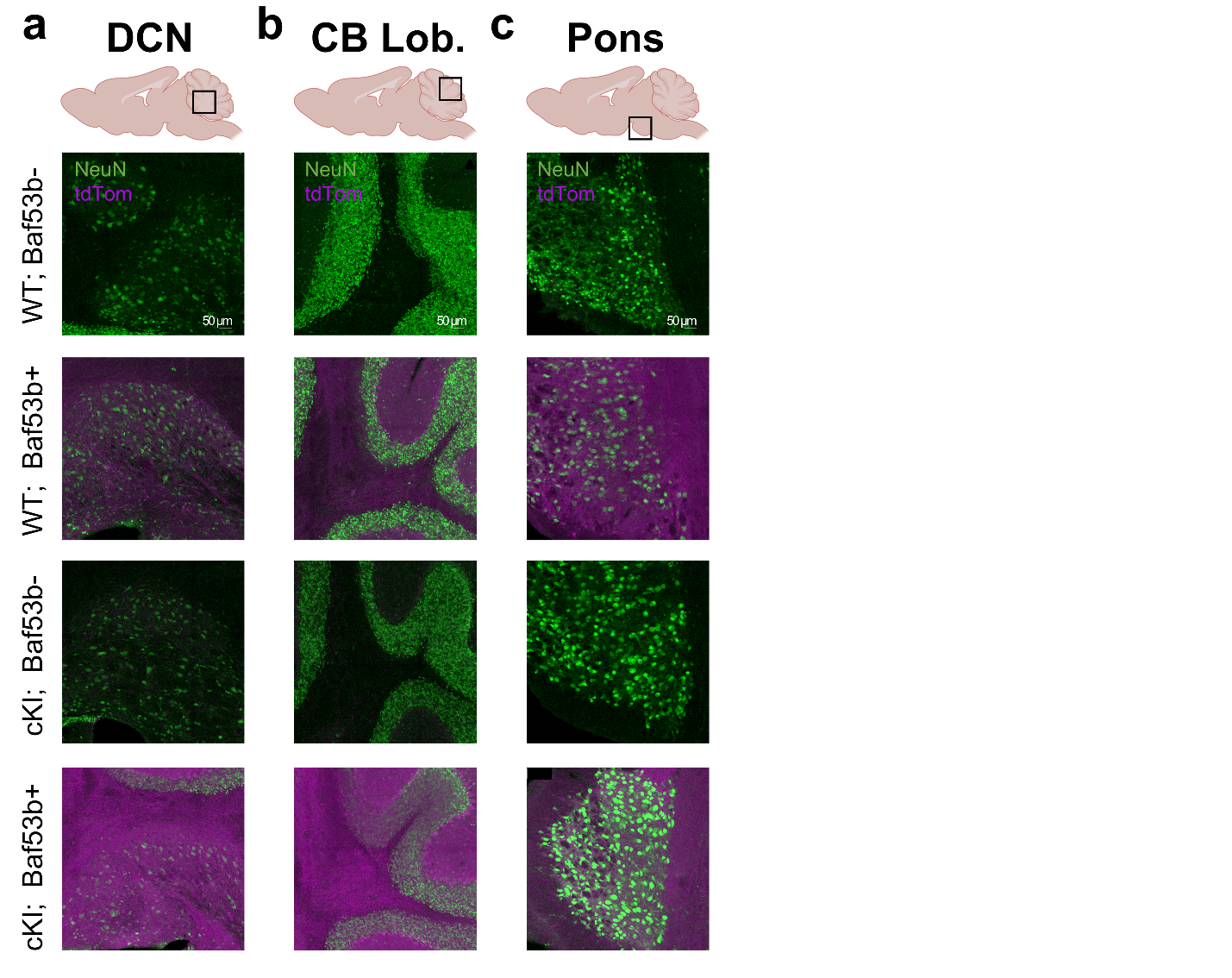
**Fig. S2 *Baf53b-Cre; tdTomato* reporter mice validate neuronal Cre recombination in SCA3-vulnerable brain regions**

Representative immunofluorescent images of 36-week-old WT; Baf53b-, WT; Baf53b+, cKI; Baf53b-, and cKI; Baf53b+ mice stained for tdTomato (tdTom) and NeuN in the **(a)** deep cerebellar nuclei (DCN, scale bar = 50 µm), **(b)** cerebellar lobules (CB Lob., scale bar = 50 µm), and **(c)** pontine nuclei (Pons, scale bar = 50 µm). All images are maximum z-projections.

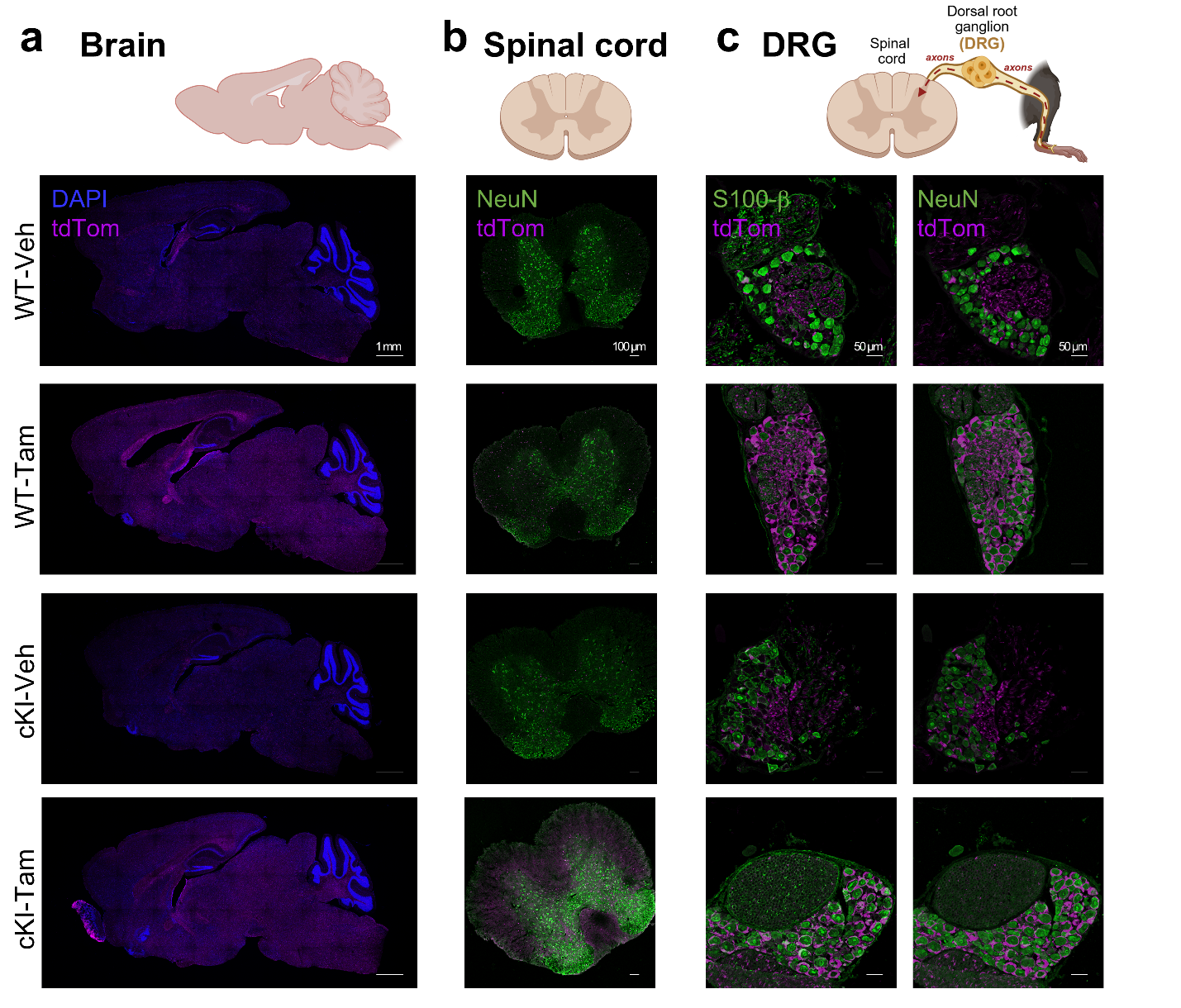
**Fig. S3 *Sox10-iCre/ERT2; tdTomato* reporter mice validate robust Cre recombination in central and peripheral nervous systems**

*Sox10-iCre/ERT2+* (iSox10) mice were injected with vehicle (Veh) or tamoxifen (Tam) at 12 weeks of age and collected at 14 weeks. **(a)** Representative immunofluorescent images of WT-Veh, WT-Tam, cKI-Veh, and cKI-Tam mice stained for tdTomato (tdTom) and DAPI in sagittal brain sections (scale bar = 1 mm). **(b)** Representative images of tdTom and NeuN staining in iSox10 spinal cord sections (scale bar = 100 µm). **(c)** Representative images of tdTom and S100-beta (left) or NeuN (right) staining in iSox10 DRG sections (scale bar = 50 µm). DRG images are a maximum z-projection, and all others are single-plane.

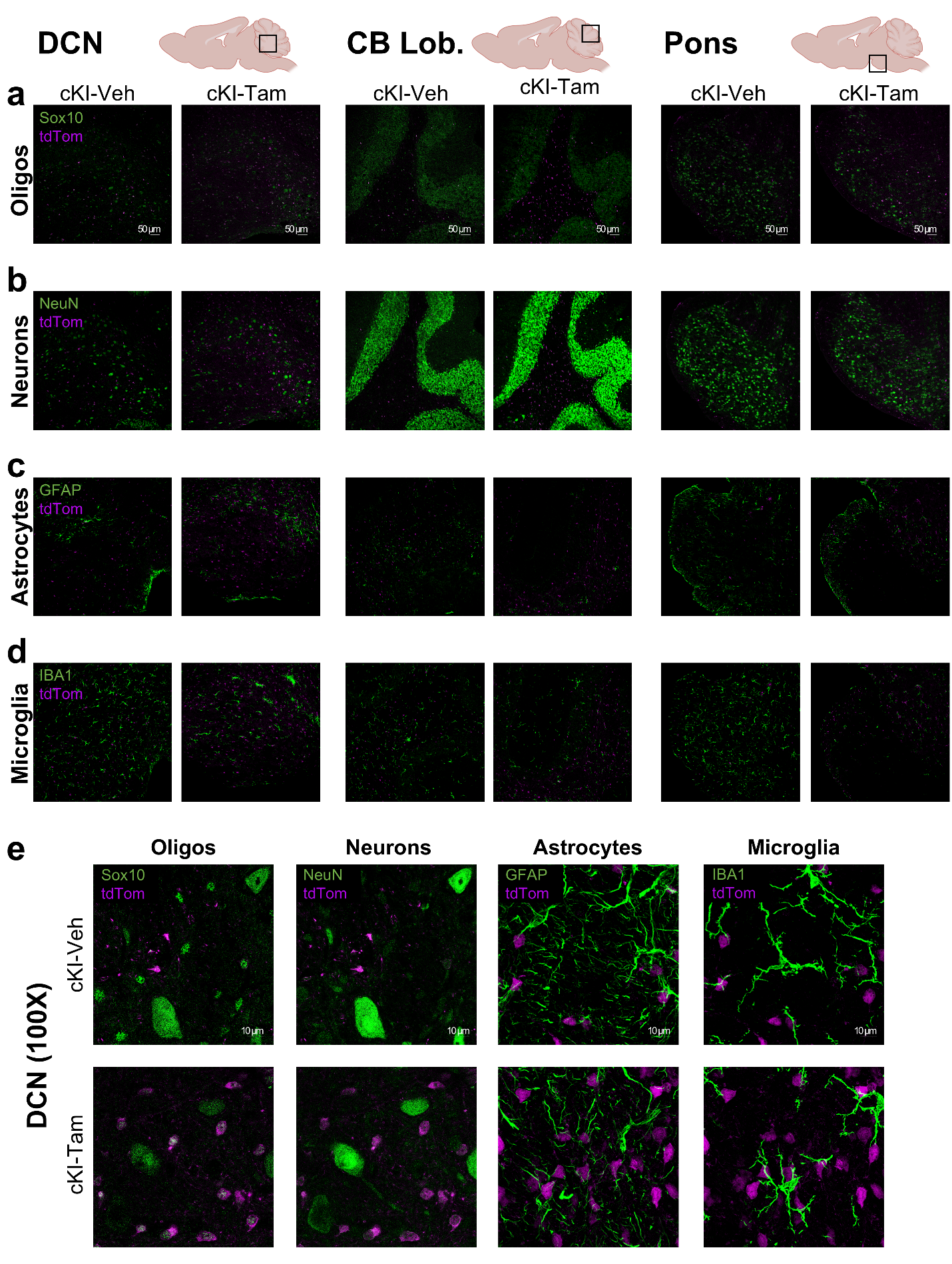
**Fig. S4 *Sox10-iCre/ERT2* reporter expression is restricted to oligodendrocytes and absent from neurons, microglia, and astrocytes in SCA3-vulnerable brain regions**

The cell type-specificity of Cre recombination was evaluated across SCA3-vulnerable brain regions of *Sox10-iCre/ERT2+* (iSox10) mice injected with vehicle (Veh) or tamoxifen (Tam) at 12 weeks of age and collected at 14 weeks. **(a)** Representative images of WT-Veh, WT-Tam, cKI-Veh, and cKI-Tam mice stained for tdTomato (tdTom) in Sox10+ oligodendrocytes (Oligos) of the deep cerebellar nuclei (DCN, left), cerebellar lobules (CB Lob., middle), and pontine nuclei (Pons, right). **(b)** Representative images of tdTom in NeuN+ neurons. **(c)** Representative images of tdTom in GFAP+ astrocytes. **(d)** Representative images of tdTom in IBA1+ microglia. Scale bar for a–d = 50 µm. **(e)** Representative high-magnification images of the cKI-Tam DCN for oligodendrocytes, neurons, astrocytes, and microglia (scale bar = 10 µm). Images of oligodendrocytes (a, e) and neurons (b, e) are single-plane, and images of astrocytes (c, e) and microglia (d, e) are maximum z-projections.

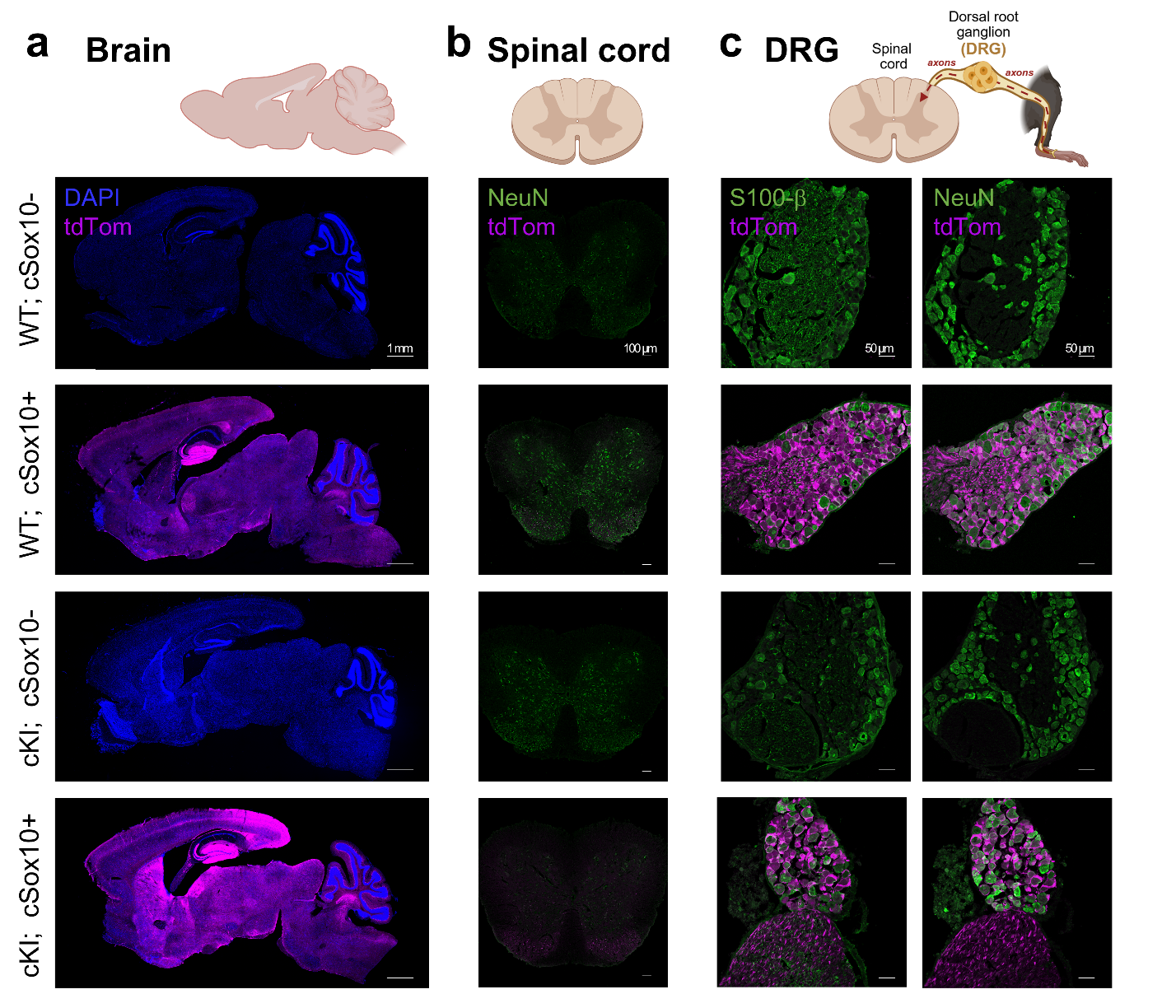
**Fig. S5 Constitutive *Sox10-Cre; tdTomato* reporter mice validate robust Cre recombination in the central and peripheral nervous systems**

**(a)** Representative immunofluorescent images of 12-week-old constitutive *Sox10-Cre* (cSox10) mice (WT; cSox10-, WT; cSox10+, cKI; cSox10-, and cKI; cSox10+) stained for tdTomato (tdTom) and DAPI in sagittal brain sections (scale bar = 1 mm). **(b)** Representative images of tdTom and NeuN staining in cSox10 spinal cord sections (scale bar = 100 µm). **(c)** Representative images of tdTom and S100-b (left) or NeuN (right) staining in cSox10 DRG sections (scale bar = 50 µm). DRG images are a maximum z-projection, and all others are single-plane.

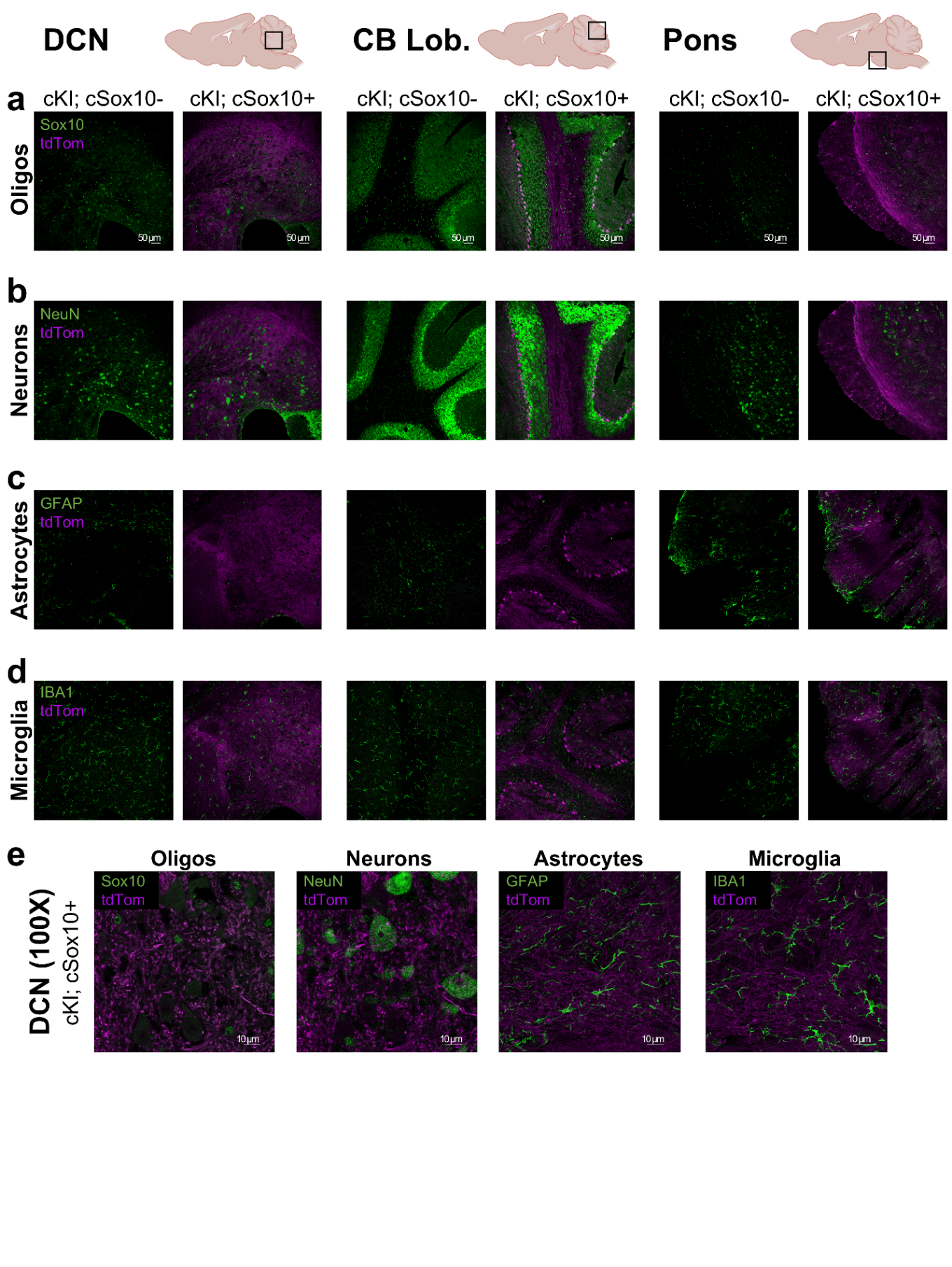

**Fig. S6 Constitutive *Sox10-Cre* strongly targets oligodendrocytes and Purkinje cells and minimally targets astrocytes and microglia in SCA3-vulnerable brain regions**

The cell type-specificity of Cre recombination was evaluated across SCA3-vulnerable brain regions in 12-week-old *Sox10-Cre* constitutive (cSox10) mice (WT; cSox10-, WT; cSox10+, cKI; cSox10-, and cKI; cSox10+). **(a)** Representative images of tdTomato (tdTom) in Sox10+ oligodendrocytes (Oligos) of the deep cerebellar nuclei (DCN, left), cerebellar lobules (CB Lob., middle), and pontine nuclei (Pons, right). **(b)** Representative images of tdTom in NeuN+ neurons. **(c)** Representative images of tdTom in GFAP+ astrocytes. **(d)** Representative images of tdTom in IBA1+ microglia. Scale bar for a–d = 50 µm. **(e)** Representative high-magnification images of the cKI; cSox10+ DCN for oligodendrocytes, neurons, astrocytes, and microglia (scale bar = 10 µm). Images of oligodendrocytes (a, e) and neurons (b, e) are single-plane, and images of astrocytes (c, e) and microglia (d, e) are maximum z-projections.

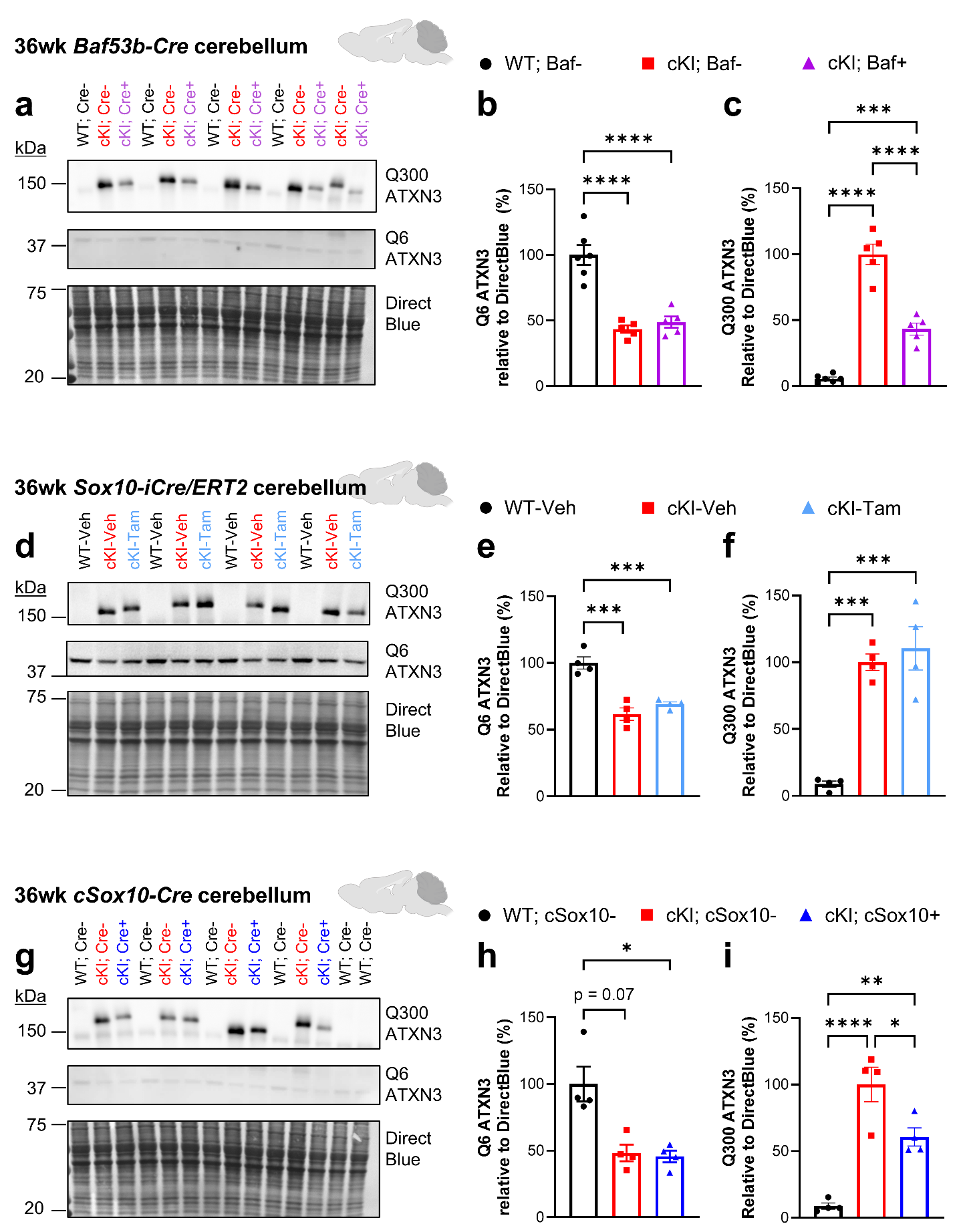
**Fig. S7 Biochemical validation of Cre target engagement in the cerebellum of late-stage *Baf53b-Cre*, *Sox10-iCre/ERT2*, and *Sox10-Cre* mice**

Representative images of Western blots for wildtype (Q6, 1H9 antibody) and mutant (Q300, 1C2 antibody) ATXN3 in the cerebellum of 36-week-old **(a)** *Baf53b-Cre* (Baf53b), **(d)** *Sox10-iCre/ERT2* (iSox10), and **(g)** *Sox10-Cre* (cSox10) mice. Quantification of wildtype (Q6) ATXN3 normalized to total protein levels measured by Direct Blue between 20 and 75 kDa in **(b)** Baf53b, **(e)** iSox10, and **(h)** cSox10 mice. Quantification of mutant (Q300) ATXN3 normalized to total protein levels in **(c)** Baf53b, **(f)** iSox10, and **(i)** cSox10 mice. n=4–6 mice/group. Graphs represent mean ± SEM, with each point as one mouse. One-way ANOVA with post-hoc Tukey’s test or nonparametric equivalent (Kruskal-Wallis with post-hoc Dunn’s test). No symbol = not significant; * p < 0.05; ** p < 0.01; *** p < 0.001; **** p < 0.0001.

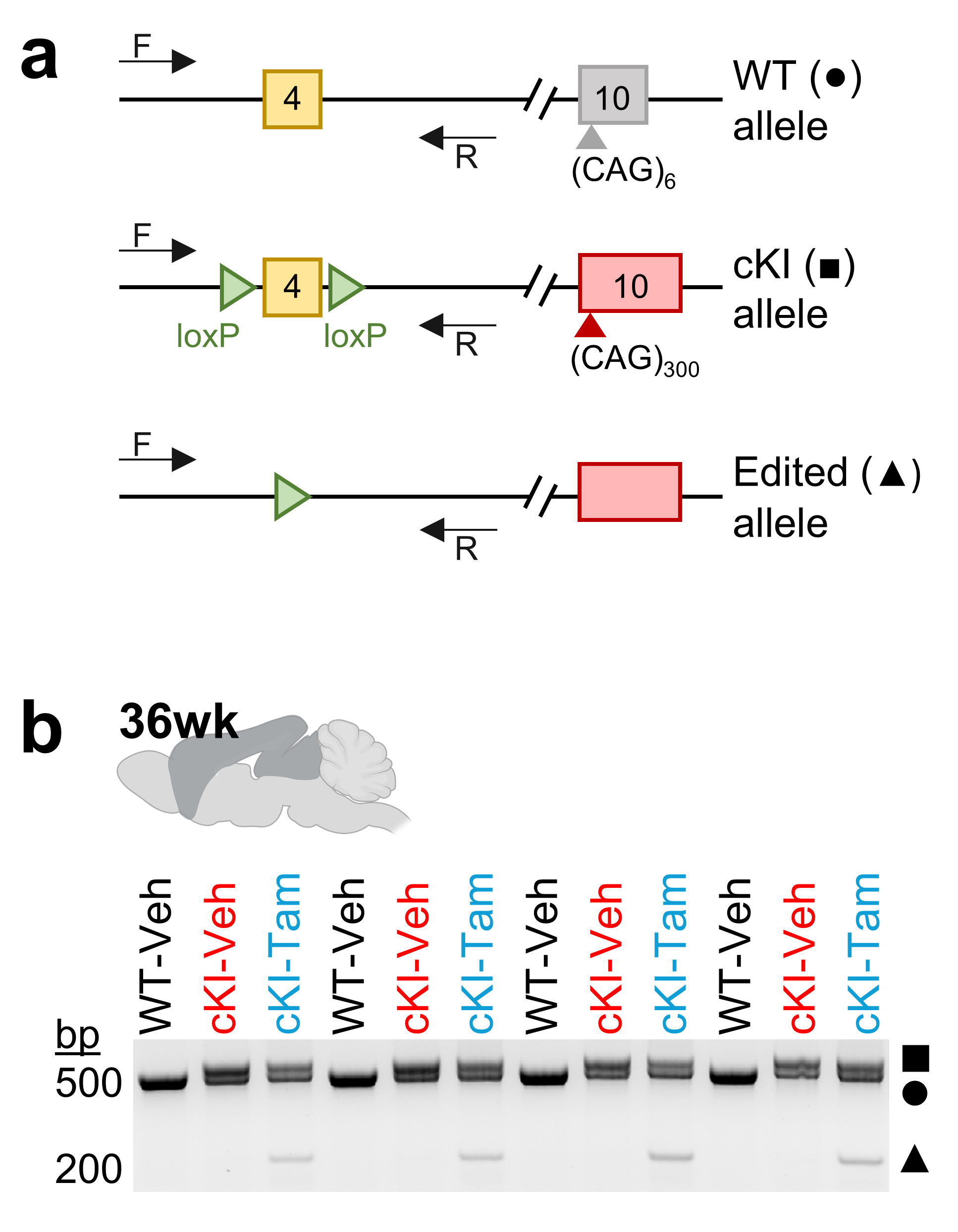

**Fig. S8 Exon 4 is excised from the mutant *Atxn3* allele in the frontal cortex of *Sox10-iCre/ERT2* mice injected with tamoxifen**

**(a)** Schematic of WT and conditional Knock-in (cKI) *Atxn3* alleles. The cKI allele contains loxP sites flanking exon 4 and a hyper-expanded 300 CAG repeats in exon 10. In Cre+ cells, exon 4 is excised, generating the edited allele. The binding sites for forward (“F”) and reverse (“R”) primers flanking exon 4 are noted. **(b)** PCR was conducted on genomic DNA isolated from the frontal cortex of 36-week-old WT and cKI mice injected with vehicle or tamoxifen at 12 weeks of age. WT-Veh mice have a single band at ~537 bp (●), representing the WT *Atxn3* allele. cKI-Veh mice have an additional band at ~605 bp (■), representing the intact cKI allele which is larger than the WT allele due to the addition of loxP sites flanking exon 4. cKI-Tam mice show bands for the intact WT and cKI alleles, as well as a smaller band at ~224 bp (▲) representing the size of the mutant *Atxn3* allele when exon 4 is excised (“edited allele”).

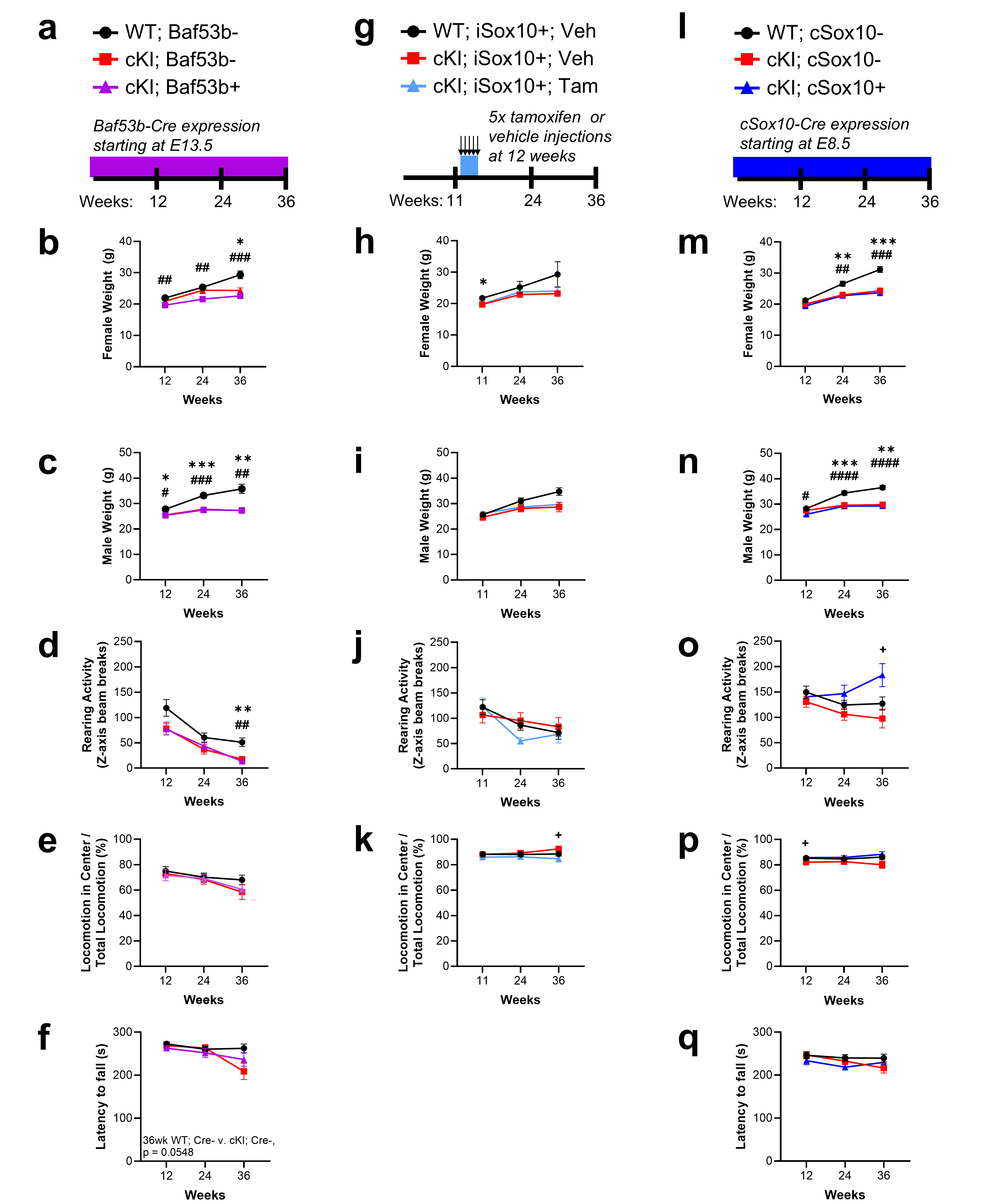
**Fig. S9 Longitudinal data for weight, rearing, anxiety-like behavior, and rotarod in *Baf53b-Cre, Sox10-Cre,* and *Sox10-iCre/ERT2* mice**

Schematic of motor behavior assessment timeline in **(a)** *Baf53b-Cre* (Baf53b), **(g)** *Sox10-iCre/ERT2* (iSox10) and **(l)** *Sox10-Cre* (cSox10) mice. **(b, h, m)** Longitudinal weight of female mice in each model (Baf53b: n=5–13 mice/group at 12wk, n=4–13 mice/group at 24wk, and n=3–10 mice/group at 36wk; iSox10: n=3–10 mice/group at 11wk, n=2–10 mice/group at 24wk, and n=2–9 mice/group at 36wk; cSox10: n=16–20 mice/group mice at 12wk, n=12–17 mice/group at 24wk, and n=5–10 mice/group at 36wk). **(c, i, n)** Longitudinal weight of male mice in each model (Baf53b: n=8–12 mice/group at 12wk, n=7–12 mice/group at 24wk, and n=7–10 mice/group at 36wk; iSox10: n=5–7 mice/group at 11wk, n=4–5 mice/group at 24wk, and n=4–5 mice/group at 36wk; cSox10: n=17–21 mice/group at 12wk, 13–17 mice/group at 24wk, and n=4–10 mice/group at 36wk). **(d, j, o)** Longitudinal rearing activity (z-axis beam breaks) on the open field apparatus in each mouse model. **(e, k, p)** Longitudinal percent of total locomotion in the center of the open field apparatus (as a readout of anxiety-like behavior) in each mouse model. **(f, q)** Latency to fall on the fourth day of the accelerated rotarod task in Baf53b and cSox10 mice, respectively. *Baf53b-Cre:* For panels d–f, n=16–24 mice/group at 12wk, 14–24 mice/group at 24wk, and 11–20 mice/group at 36wk. *Sox10-iCre/ERT2*: For panels j and k: n=9–15 mice/group at 11wk, n=6–15 mice/group at 24wk, and n=6–13 mice/group at 36wk. *Sox10-Cre:* For panels o–q, n=27–41 mice/group at 12wk, n=26–34 mice/group at 24wk, and n=13–19 mice/group at 36wk. Graphs display mean ± SEM. Mixed-effects analysis with Tukey’s multiple comparisons test. * indicates the WT; Cre- vs. KI; Cre- comparison for the *Baf53b-Cre* and *cSox10-Cre* models, and WT; iSox10+; Veh vs. cKI; iSox10+; Veh for the *Sox10-iCre/ERT2* model. # indicates the WT; Cre- vs. KI; Cre+ comparison for the *Baf53b-Cre* and *cSox10-Cre* models, and WT; iSox10+; Veh vs. cKI; iSox10+; Tam for the *Sox10-iCre/ERT2* model. + indicates the cKI; Cre+ vs. cKI; Cre- comparison for the *Baf53b-Cre* and *cSox10-Cre* models, and cKI; iSox10+; Tam vs. cKI; iSox10+; Veh comparison for the *Sox10-iCre/ERT2* model. No symbol = not significant; one symbol, p < 0.05; two symbols, p < 0.01; three symbols, p < 0.001; four symbols, p < 0.0001.

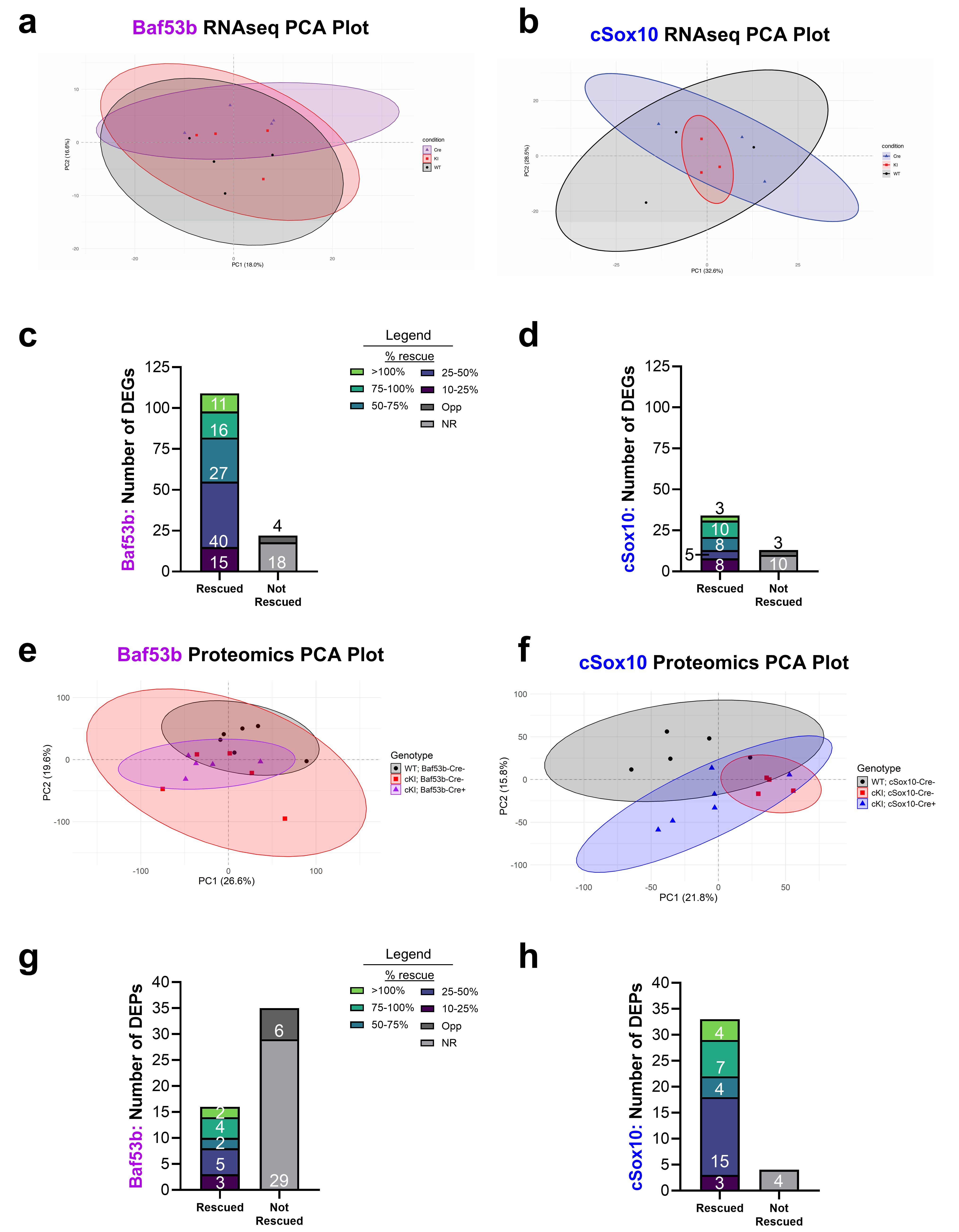
**Fig. S10 Principal component analysis and rescue plots for *Baf53b-* and *Sox10-Cre* cerebellar bulk RNA sequencing and proteomics data**

Principal component analysis (PCA) plots for cerebellar bulk RNA sequencing (n=3–4 mice/group) on 36-week-old **(a)** *Baf53b-Cre* (Baf53b) and **(b)** *Sox10-Cre* (cSox10) mice. **(c)** Plot denoting the distribution of differentially expressed genes (DEGs) in cKI; Cre- mice relative to WT; Cre- controls that are rescued by Baf53b-Cre. **(d)** Rescue plot for cSox10 DEGs. PCA plots for cerebellar proteomics (n=4–6 mice/group) on 36-week-old **(e)** Baf53b and **(f)** cSox10 mice. **(g)** Rescue plot for Baf53b differentially expressed proteins (DEPs). **(h)** Rescue plot for cSox10 DEPs. For PCA plots, each point is one mouse, with a 95% confidence ellipse computed for each group. PC1 = Principal Component 1, with percent explained variance in parentheses. PC2 = Principal Component 2. Rescue was defined as a gene or protein with a minimum 10% change in cKI; Cre+ mice relative to cKI; Cre- mice, in the direction of WT; Cre-. NR indicates not rescued gene or proteins, and Opp indicates those changed in the opposite direction of WT.

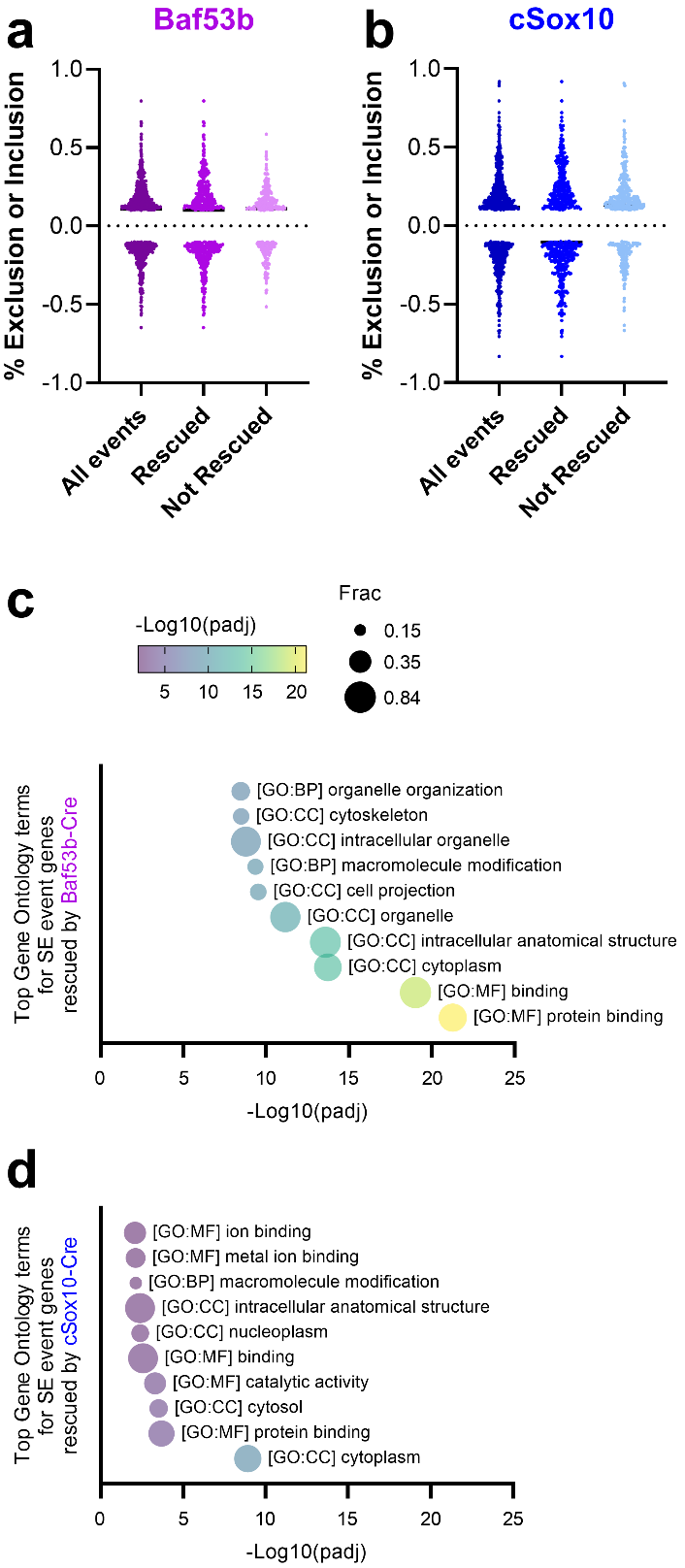

**Fig. S11 Percent exclusion versus inclusion and functional enrichment analysis for *Baf53b*- and *Sox10-Cre* significant skipped exon events**

Plot of delta PSI (percent spliced in) for the skipped exon events that were significant in the comparison of cKI; Cre- mice relative to WT; Cre- mice, displaying the distribution for all events, rescued events, and not rescued events in the **(a)** *Baf53b-Cre* (Baf53b) and **(b)** *Sox10-Cre* (cSox10) 36-week cerebellar datasets, with an even distribution between exon inclusion (above 0) and exclusion (below 0). Each point is one event, with n=1029 significant events in Baf53b and n=1138 significant events in cSox10. **(c)** Top ten statistically significant Gene Ontology (GO) terms for the SE event genes rescued by Baf53b. Frac represents the fraction of genes in the GO term contained in the dataset. **(d)** Top GO terms for the SE event genes rescued by cSox10. n=3–4 mice/group.
